# Increased Enhancer—Promoter Interactions during Developmental Enhancer Activation in Mammals

**DOI:** 10.1101/2022.11.18.516017

**Authors:** Zhuoxin Chen, Valentina Snetkova, Grace Bower, Sandra Jacinto, Benjamin Clock, Atrin Dizehchi, Iros Barozzi, Brandon J. Mannion, Ana Alcaina-Caro, Javier Lopez-Rios, Diane E. Dickel, Axel Visel, Len A. Pennacchio, Evgeny Z. Kvon

## Abstract

Remote enhancers are thought to interact with their target promoters via physical proximity, yet the importance of this proximity for enhancer function remains unclear. Here, we investigate the 3D conformation of enhancers during mammalian development by generating high-resolution tissue-resolved contact maps for nearly a thousand enhancers with characterized *in vivo* activities in ten murine embryonic tissues. 61% of developmental enhancers bypass their neighboring genes, which are often marked by promoter CpG methylation. The majority of enhancers display tissue-specific 3D conformations, and both enhancer–promoter and enhancer–enhancer interactions are moderately but consistently increased upon enhancer activation *in vivo*. Less than 14% of enhancer–promoter interactions form stably across tissues; however, these invariant interactions form in the absence of the enhancer and are likely mediated by adjacent CTCF binding. Our results highlight the general significance of enhancer– promoter physical proximity for developmental gene activation in mammals.

## Introduction

Enhancers, or *cis*-regulatory elements, ensure precise spatiotemporal control of gene expression during development. This process is mediated by transcription factors (TFs) and co-activators, which relay regulatory information from enhancers to their target promoters, across distances that can exceed one megabase^1–4^. This enhancer–promoter (E–P) communication is thought to occur within so-called topologically associated domains (TADs), fundamental organizational units of the genome formed through the process of loop extrusion by cohesin and CCCTC-Binding Factor (CTCF)^5–7^. Disruption of TADs or intra-TAD chromatin interactions can cause erroneous downregulation of gene expression or gene activation and can lead to human disease, indicating the importance of proper E–P communication for gene activation^8–10^.

Remote enhancers are thought to communicate with their target genes via physical proximity established by chromatin looping^11–14^. However, whether physical proximity is linked to enhancer function remains unclear. One model suggests that E–P contacts are formed only during gene activation. Indeed, the establishment of E–P interactions at many genetic loci occurs coordinately with gene transcription^15–18^. In line with this, artificial tethering of an enhancer to the developmentally silenced β-globin promoter results in an ectopic gene activation^19^, suggesting a potentially instructive role of chromatin looping in E–P communication and gene activation. An alternative model is that E–P contacts are stable and/or pre-formed and thus not temporally linked to gene activation. For example, mouse limb enhancers at the *HoxD* and *Shh loci*, human fibroblast and keratinocyte enhancers, and many early *Drosophila* enhancers appear to form E–P chromatin loops even when the genes are not expressed^18,20–23^. In a third model, there is no association between gene activation and E–P physical proximity^24^, and in some cases, an increase in E—P distance is observed upon gene activation, challenging a simple looping model^25,26^. While all these models exist in principle, the predominant mode of activation for *bona fide* developmental enhancers remains unclear since past research has focused on well-studied genetic loci or enhancers defined based on the presence of open chromatin, co-activators, eRNAs, or enhancer-associated histone modifications, thus making it challenging to separate functional E—P interactions from other types of chromatin interactions^27^.

To better understand E–P interactions during mammalian development, we utilized a unique resource of experimentally verified human and mouse enhancers^28^. Many of these enhancers have been shown to be critical for developmental and disease processes^8,29–33^. However, the 3D nuclear organization of these loci remains largely uncharacterized. We thus generated high-resolution enhancer interactome maps across 10 mouse embryonic tissues for 935 *bona fide* developmental enhancers with characterized *in vivo* activity at mid-gestation. We identified thousands of enhancer contacts and found that most enhancer loci display tissue-specific 3D conformations. Moreover, developmental enhancers display higher interaction frequencies with promoters and neighboring enhancers in tissues where they are active. We also show that invariant E—P interactions are less prevalent and likely form independently of enhancer activity. 61% of developmental enhancers skip their immediate neighboring genes, which are often marked by promoter DNA methylation. Our results provide a global view of tissue-specific enhancer 3D chromatin conformation and support the broad importance of E–P physical proximity for developmental gene activation.

## Main text

### Enhancer interactome for 935 developmental enhancers across 10 embryonic tissues

To create a map of *in vivo* enhancer-centric chromatin interactions in developing mouse embryos, we used the VISTA Enhancer Browser, a unique resource of human and mouse enhancers with *in vivo* activities experimentally validated in transgenic mice^28^. This resource verifies, and thus allows direct comparison of, tissue/cell types in which each tested enhancer is active or inactive. We created a sizable and robust core set of experimentally verified *in vivo* enhancers comprising 935 enhancers with highly reproducible activities in mouse embryonic tissues at mid-gestation (embryonic day 11.5). Tissues in which enhancers were active included the forebrain, midbrain, hindbrain, neural tube, craniofacial structures, limb buds, heart and other tissues and cell types (see **Supplementary Table 1**). To assess tissue-specific chromatin interactions centered on these enhancers, we collected 10 tissues from E11.5 mouse embryos (forebrain, midbrain, hindbrain, neural tube, face, forelimb, hindlimb, heart, tail and trunk) with two biological replicates per tissue and performed the enhancer capture Hi-C (Fig. 1a and Methods). This diverse tissue panel represents all major embryonic organs in which selected enhancers are active and for which extensive chromatin state maps were created as part of the ENCODE project^34^. We designed RNA probes (Agilent SureSelect platform) targeting each of the 935 enhancers, as well as 176 promoters and 87 elements with no reproducible enhancer activity at E11.5 as negative controls (Fig. 1a, **Methods** and **Supplementary Table 1**).

**Fig. 1:**
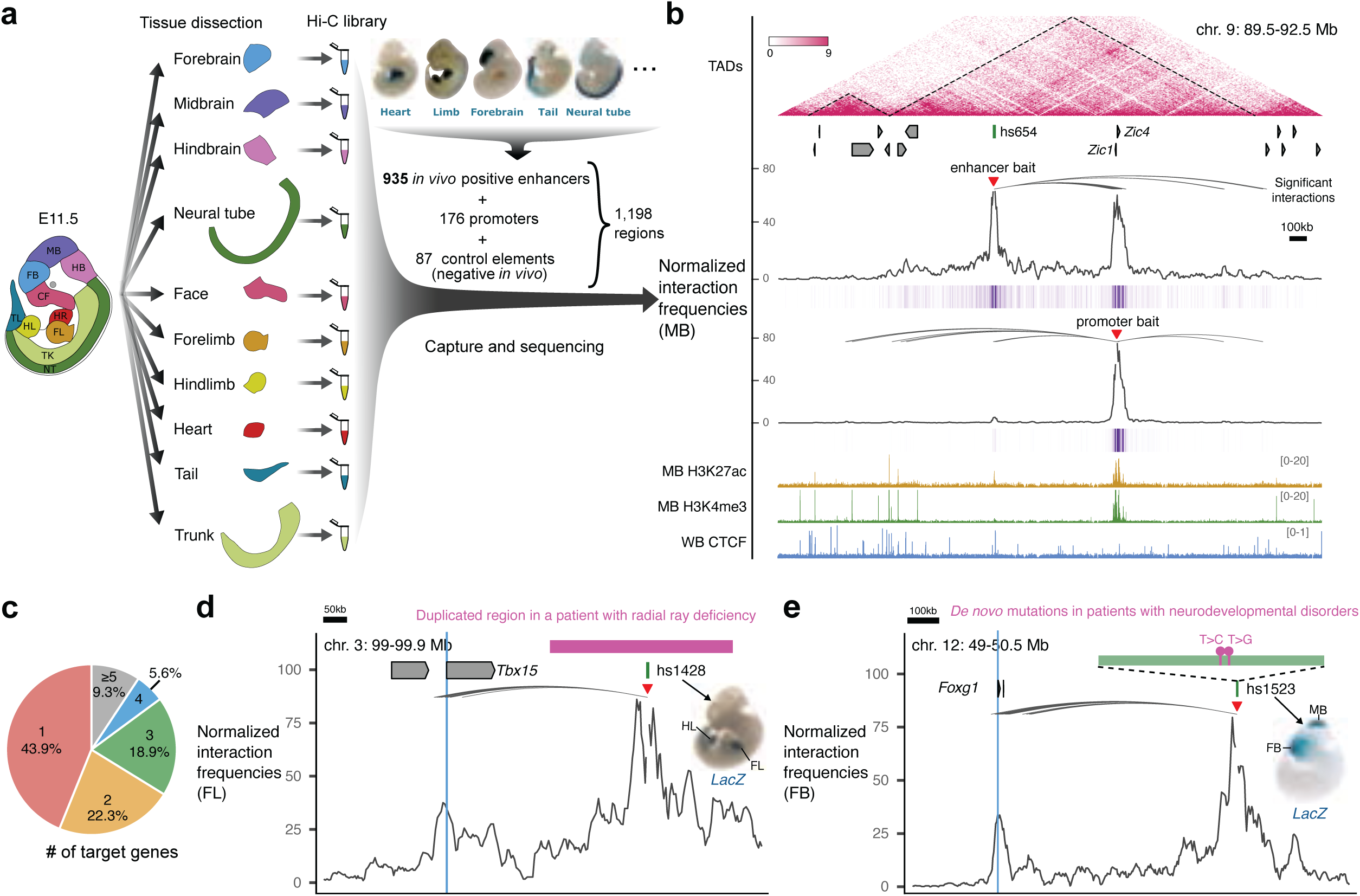
Identification of enhancer-centric chromatin interactions in 10 mouse embryonic tissues. **a**, Experimental design. Ten tissue samples from E11.5 mouse embryos were used to prepare Hi-C libraries followed by oligonucleotide capture with probes targeting 1,198 baited regions, including 935 enhancers (representative enhancer activities are shown above), 176 promoters and 87 control elements. **b**, Enhancer capture Hi-C identifies chromatin interactions of enhancers. A 3 Mb region containing the hs654 midbrain enhancer (chr9:89500000-92500000; mm10) is shown with the following annotations from top to bottom: TADs (dashed lines outline TAD boundaries)^74,75^; Refseq genes; normalized hs654-centered chromatin interaction frequencies in midbrain (MB) shown as plot and purple heat map below; normalized *Zic1/Zic4*-promoter-centered chromatin interaction frequencies; H3K27ac and H3K4me3 ChIP-seq profiles in midbrain at E11.5; CTCF ChIP-seq profile in whole brain (WB) at E12.5^34,76,77^. The average bin size is ∼3kb. Curved lines indicate significant interactions. **c**, Pie chart showing the percentage of enhancers interacting with different number of genes. **d**, The hs1428 limb enhancer (green box) is in a non-coding region (purple bar) which is duplicated in patients with radial ray deficiency (pink box indicates homologous region in the mouse genome). The hs1428 limb enhancer forms significant chromatin interactions with the promoter of *Tbx15* (highlighted in blue) located ∼400 kb away (chr3:99,000,000-99,900,000; mm10)^78^ in the forelimb (FL). **e**, Two de novo rare variants (purple boxes) identified in patients with neurodevelopmental disorders^79,80^ are in the hs1523 (green bar) forebrain/midbrain enhancer which forms strong significant interactions with the promoter of *Foxg1* (highlighted in blue) located ∼700 kb away (chr12:49,121,092-50,469,462; mm10) in the forebrain (FB). Red arrowheads indicate capture Hi-C viewpoints.

After restriction fragment pooling and quality control we identified a total of 24,657 significant interactions across all tissues, 17,988 of which were baited on enhancers. ∼80% of enhancer-centric interactions were called within the same TAD (**Extended Data Fig. 1a-d, **Supplementary Table 2**** and **Methods**). These interactions included E–P (2,818), enhancer–enhancer (E–E) (5,612), enhancer–CTCF (5,140) and other types of contacts (Extended Data Fig. 1d). Most enhancers only interacted with one or two genes with a median distance between an enhancer and a target promoter of ∼410 kb (Fig. 1c and Extended Data Fig. 1f). For example, in the midbrain, the hs654 enhancer displayed the strongest significant interaction with promoters of two adjacent genes, *Zic1* and *Zic4*, located ∼600 kb away. Reciprocally the viewpoint containing the *Zic1* and *Zic4* promoters (located ∼3 kb from each other) also showed significant interaction with the hs654 enhancer (Fig. 1b).

To provide orthogonal support for the functional relevance of identified chromatin interactions we compared them with ENCODE chromatin data that was generated for an overlapping set of tissues from E11.5 mouse embryos. We found that the 935 *in vivo* positive enhancers and 176 promoters contacted other elements annotated by ENCODE (promoters, enhancers, CTCF sites) significantly more often than the negative 87 control regions, thus supporting the enhancer interactions identified above (Extended Data Fig. 1g, h).

We also identified significant tissue-specific chromatin interactions between enhancers overlapping mutations implicated in human congenital disorders and their putative target genes in relevant tissues. These examples included previously characterized enhancers involved in congenital malformations and autism as well as enhancer variants identified in patients with neurodevelopmental disorders with previously unknown regulatory targets (Fig. 1d,e, Extended Data Fig. 2 and **Supplementary Table 3**). These results provide additional evidence for the specific regulatory connection between disease-associated enhancers and their *in vivo* target genes and further support E–P chromatin interactions identified by capture Hi-C.

### Most enhancers bypass adjacent genes, which are often methylated

Nearly 61% of enhancers in our study did not interact with the promoters of adjacent genes but instead contacted more distal genes (Fig. 2a). For example, the hs271 forebrain enhancer strongly interacts with the promoter of *Nrf21* located ∼650 kb away but does not form any significant interactions with the more proximally located *Pou5f2* promoter (Fig. 2b,c). Similarly, a cluster of three forebrain enhancers, hs267, hs266 and hs853, interacted with the *mir9-2* promoter located ∼800 kb away, skipping over the more proximal Tmem161b promoter (Extended Data Fig. 3a).

**Fig. 2:**
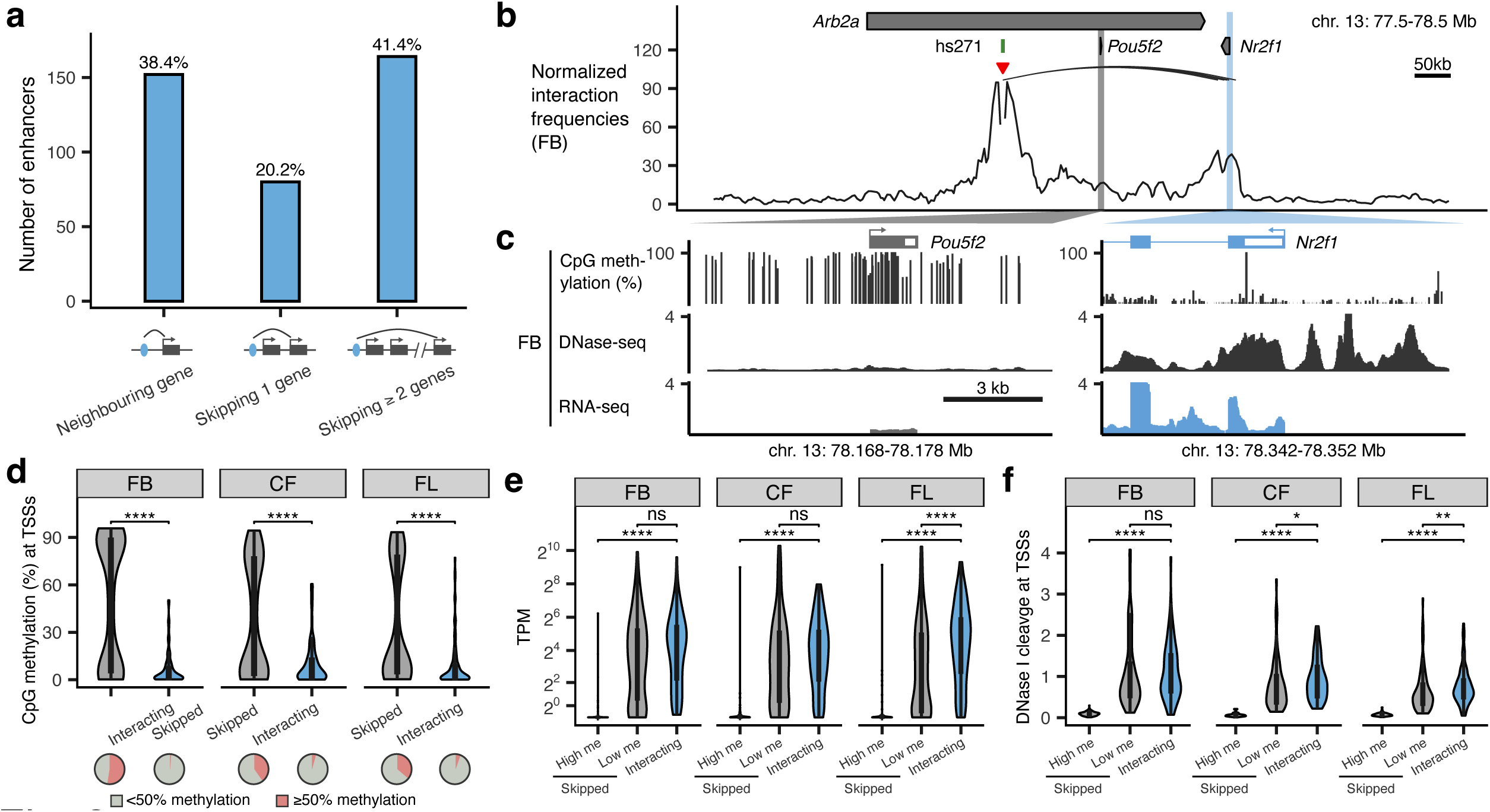
Properties of promoters that are skipped by remote enhancers. **a**, Barplot showing enhancers grouped by their genomic positions relative to the interacting genes. Diagram below shows corresponding schematic gene loci in which enhancer (blue oval) interacts with a neighboring gene (left), skips one gene (middle) or skips two or more genes (right). Arches indicate significant interactions. **b**, Normalized capture Hi-C data from the viewpoint of the hs271 enhancer (red arrowhead) is shown with significant interactions (black arches) in the forebrain at E11.5 (chr13:77,500,000-78,500,000; mm10). *Pou5f2* and *Nr2f1* promoters are highlighted in grey and blue. **c**, CpG methylation, DNase-seq and RNA-seq profiles at *Pou5f2* and *Nr2f1* promoters in E11.5 forebrain^34,76,81^. **d-f**, The CpG methylation (**d**), mRNA expression levels (**e**, transcript per million (TPM)) and DNase signal (**f**) of enhancer-interacting and skipped promoters in tissues where enhancers are active (FB, forebrain; CF, face; FL, forelimb). The number of skipped and interacting promoters in panel **d** are *n*=265 and *n*=90 (FB), *n*=144 and *n*=71 (CF) and *n*=182 and *n*=96 (FL) and the *P* values are 3.6×10^-17^, 3.9×10^-07^, 2.2×10^-11^, respectively. The number of high and low methylated skipped as well as interacting promoters in panel **e** are *n*=134, *n*=121 and *n*=90 (FB), *n*=56, *n*=81 and *n*=71 (CF) and *n*=64, *n*=111 and *n*=96 (FL) and the *P* values are 1×10^-35^, 1.3×10^-18^, 6.9×10^-22^ and 6.4×10^-5^, respectively. The number of high and low methylated skipped as well as interacting promoters in panel **f** are *n*=139, *n*=126 and *n*=90 (FB), *n*=58, *n*=86 and *n*=71 (CF) and *n*=66, *n*=116 and *n*=96 (FL) and the *P* values are 2.4×10^-34^, 2.9×10^-22^ and 0.012, 7.8×10^-25^ and 0.0039, respectively. High me / Low me, high / low methylation at skipped promoters (≥50% or <50% CpG methylation within ± 1 kb from TSS). *P*-values were calculated using the two-sided Wilcoxon rank test and adjusted for multiple testing. For the boxplots in panels d-f, the central horizontal lines are the median, with the boxes extending from the 25th to the 75th percentiles. The whiskers further extend by ±1.5 times the interquartile range from the limits of each box.

All skipped genes could be divided into two categories based on their epigenetic status (Fig. 2d and Extended Data Fig. 4). For example, in the forebrain, 52.4% of skipped genes were methylated and not accessible at their promoters (80.8% average CpG methylation at TSSs; 8-fold lower DNA accessibility than interacting genes, *P* < 0.0001; Fig. 2d,f) and displayed 56-fold lower expression levels than interacting genes (P < 0.0001; Fig. 2e). On the other hand, 47.6% of skipped genes in the forebrain were demethylated and accessible at their promoters similarly to promoters of interacting genes (Fig. 2d,f). These genes displayed expression levels comparable to interacting genes (Fig. 2e). We observed the same trends in all seven tissues for which matched expression and epigenomic data was available (Fig. 2d-f and Extended Data Fig. 4).

Interestingly, promoters of skipped genes did not display significantly higher levels of trimethylation at histone H3 lysine 27 (H3K27me3) or lysine 9 (H3K9me3) (Extended Data Fig. 4d,e), indicating that polycomb silencing and heterochromatin may not play a major role in regulating E–P selectivity. Taken together, our data indicate that most developmental enhancers in our study bypass neighboring genes, which are often inactive and marked by promoter CpG methylation.

### Enhancer knock-outs validate E–P chromatin interactions

To assess the functionality and specificity of identified E–P chromatin interactions, we created knock-out mice for hs654, hs267, hs266 and hs853 brain enhancers (Fig. 3 and Extended Data Fig. 5). All four enhancers form significant chromatin interactions with promoters of their putative target genes in the mouse embryonic brain at E11.5 (*Zic1/Zic4* for hs654 and *mir9-2* for hs267, hs266 and hs853; Figs. 1b, 4b and Extended Data Fig. 3a). We created two mouse knock-out lines, one carrying a deletion of hs654 (Δ*hs654*) and the other carrying a deletion of the hs267/hs266/hs853 enhancers (Δ*hs267/hs266/hs853*) and assessed tissues specific gene expression by RNA-seq (Extended Data Fig. 5). In Δhs654/Δhs654 mice, Zic4 RNA expression in the midbrain is reduced by ∼34% compared with wild-type levels (P_adj_ < 9.5 × 10^−3^, Fig. 3c) supporting the functional relevance of the hs654-Zic4 chromatin interaction in embryonic midbrain. Zic1 expression was reduced by ∼18%, albeit not statistically significant, and no other genes were significantly down- or upregulated in Δhs654/Δhs654 mice (Fig. 3c). Mice homozygous for the hs267/hs266/hs853 deletion show downregulation of *C130071C03Rik* (*mir9-2* precursor transcript) by ∼64% compared with the wild-type (*P_adj_* < 7.8 × 10^−32^, Fig. 3d). Notably there was no significant change in *Tmem161b* expression or any other gene in *cis*, indicating that these three enhancers specifically control the expression of *mir9-2* as predicted by chromatin interactions between hs267/hs266/hs853 and the *mir9-2* promoter but not the *Tmem161b* promoter (Extended Data Fig. 3a). Overall, the loss of enhancers results in a large decrease in transcription of interacting target genes, which supports that E–P chromatin interactions identified by enhancer capture Hi-C are functional and specific.

**Fig. 3:**
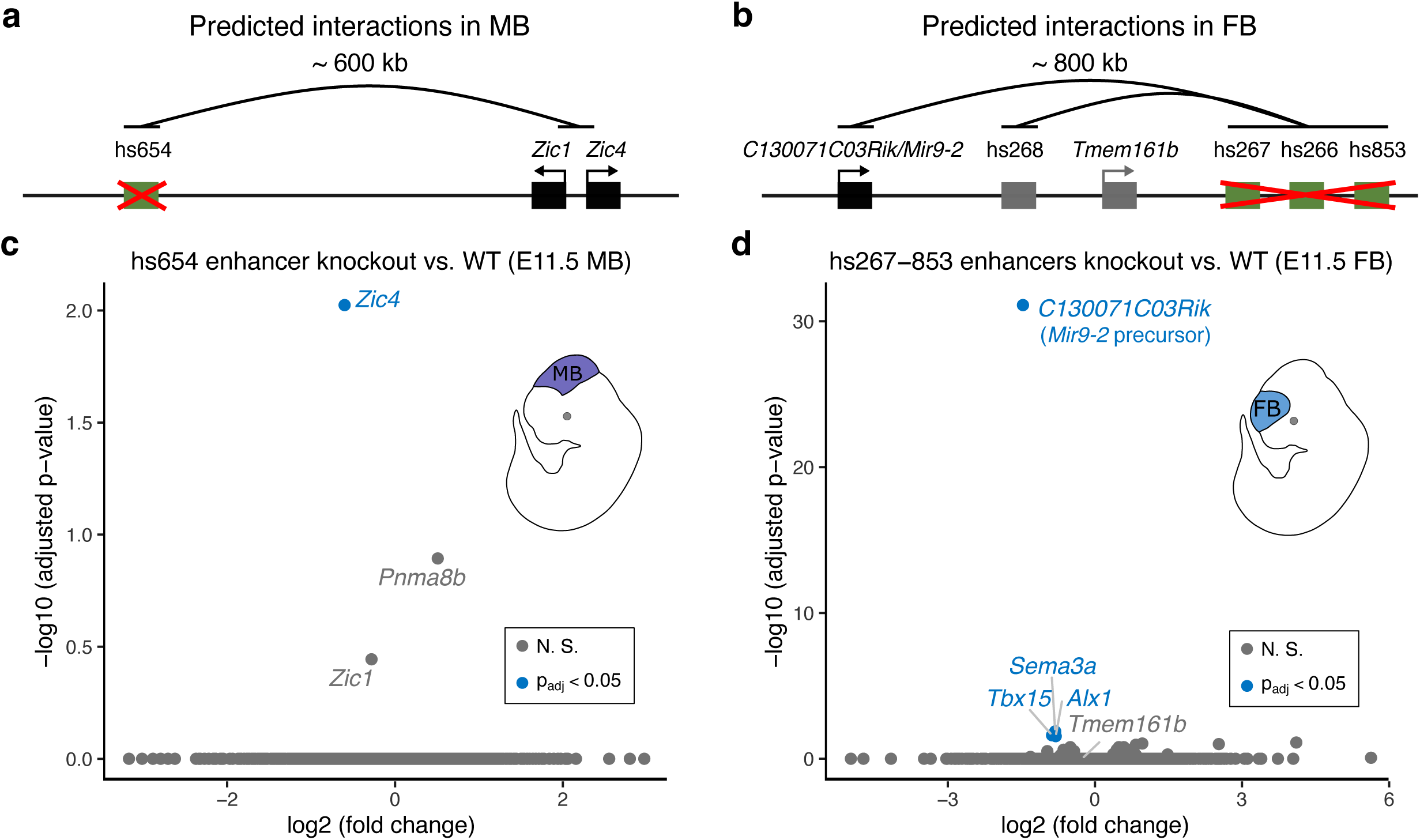
Enhancers are required for the expression of interacting genes. Knock-out analysis of hs654 and hs267/hs266/hs853 enhancers. **a,b**, Predicted chromatin interactions (black arches) between enhancers (green boxes) and target genes (black boxes) are shown. Gene and enhancer models are not drawn to scale. **c,d**, Transcriptome-wide mRNA expression changes in E11.5 whole midbrain (MB) of hs654 knock-out mice (**c**) and in E11.5 forebrain (FB) of hs267/hs266/hs853 knockout mice (**d**) relative to wildtype mice (WT). Points indicate individual genes, with blue indicating statistically significant differences after adjustment for multiple comparisons (*P*_adj_ < 0.05). N. S., not significant. *P* values were calculated using DESeq2.

**Fig. 4:**
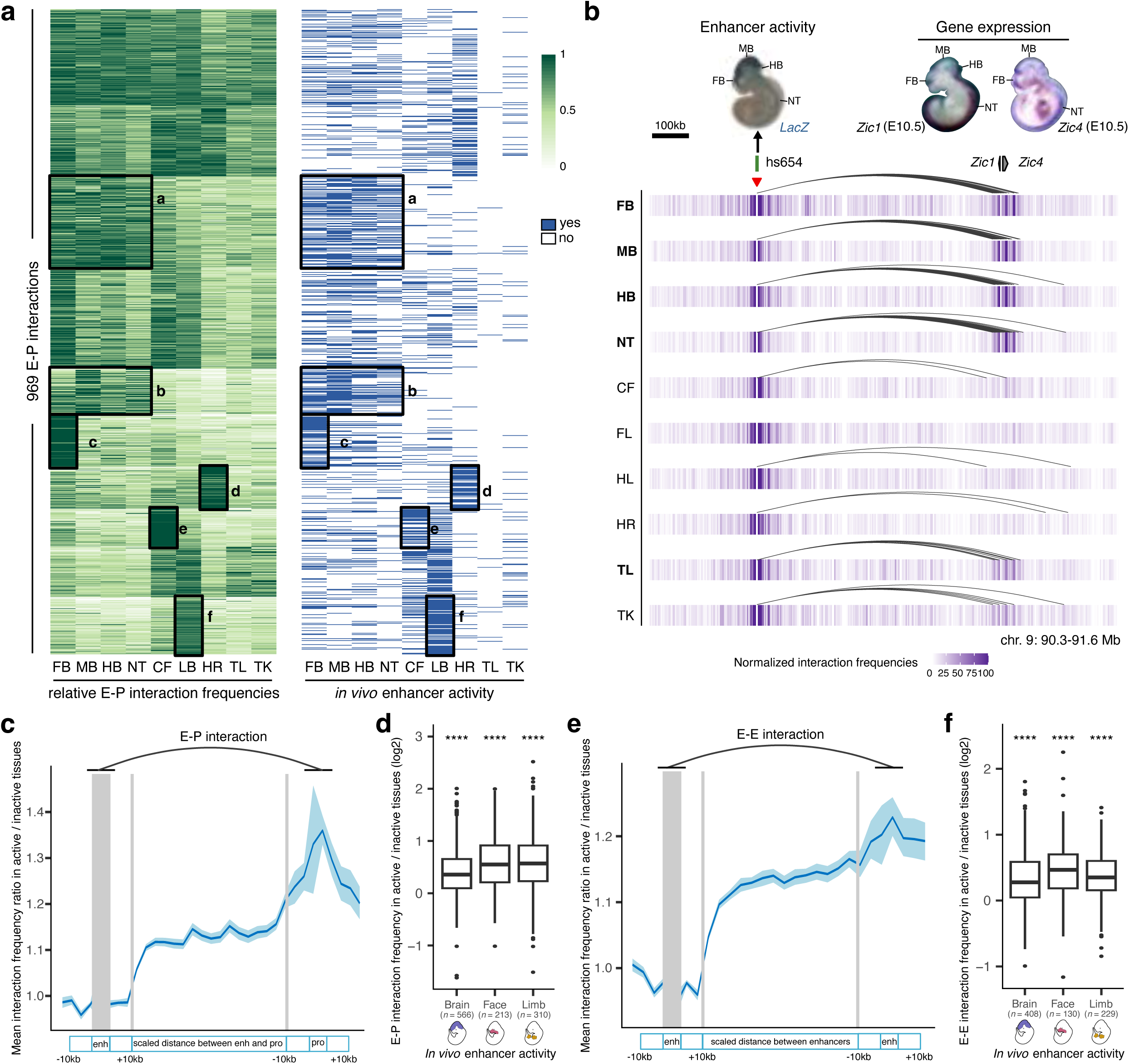
Tissue specificity of developmental enhancer interactions. **a**, Heatmap showing relative E–P chromatin interaction frequencies (scaled to the max value among tissues in each E–P interaction, green) and the *in vivo* enhancer activities (blue) of 969 E–P chromatin interactions. k-means clustering (*k* = 10) was performed on interaction frequencies. The six highlighted tissue-specific interaction clusters match *in vivo* enhancer activities. **b**, Interaction profiles across 10 tissues centered on the hs654 enhancer (red arrowhead indicates capture Hi-C viewpoint). The top left shows hs654 enhancer activity in a transgenic mid-gestation (E11.5) mouse embryo. Top right images show *Zic1* and *Zic4* mRNA whole-mount *in situ* hybridization (WISH) at E10.5 (Images reproduced with permission from Gene Expression Database (GXD; *Zic4*)^35^ and Embrys database (http://embrys.jp; *Zic1*). Heatmaps with normalized interaction frequencies in each of the 10 tissues are shown below. Curved lines indicate significant interactions. **c, e**, Average ratio of E–P or E–E interaction frequency between active and inactive tissues based on the analysis of 946 E–P or 640 E–E chromatin interactions are shown (see **Methods** for details of normalization procedure). Light blue shading indicates 95% confidence intervals estimated by non-parametric bootstrapping. **d, f**, Average ratio of E–P or E–E interaction frequency between active and inactive tissues for enhancers active in brain, face and limb (see **Extended Data Fig. 6a** and **Extended Data Fig. 8b** for other tissues). The *P* values for E–P interactions are 5.07×10^-61^ (Brain), 6.1×10^-28^ (Face), 6.21×10^-43^ (Limb). The *P* values for E–E interactions are 3.3×10^-38^ (Brain), 1×10^-17^ (Face), 1.5×10^-29^ (Limb). FB, forebrain. MB, midbrain. HB, hindbrain. CF, craniofacial mesenchyme. HR, heart. FL, forelimb. HL, hindlimb. TK, trunk. TL, tail. NT, neural tube. For the boxplots in panels d and f, the central horizontal lines are the median, with the boxes extending from the 25th to the 75th percentiles. The whiskers further extend by ±1.5 times the interquartile range from the limits of each box.

### Enhancer interactions are more frequent when enhancers are active *in vivo*

The general extent to which E–P interaction frequency correlates with *in vivo* enhancer activity at most developmental loci is unclear yet critical for understanding the spatio-temporal control of long-range gene regulation during development. To address this, we systematically compared tissue-specific enhancer activities with corresponding E–P interactions in different parts of the embryo. We selected 969 interacting E–P pairs identified by enhancer capture Hi-C where gene expression matched enhancer activity in at least one tissue (**Supplementary Table 2** and **Methods**). We then systematically examined E–P chromatin interaction profiles in each of the ten tissues and compared them with the experimentally determined *in vivo* activities of corresponding enhancers in each of these tissues. Clustering of 969 E–P interactions across ten tissues revealed a strong correlation with *in vivo* enhancer activities (logistic regression, *P* = 9.7 × 10^−46^, Fig. 4a and Extended Data Fig. 6b). Enhancers active in the central nervous system displayed higher interaction frequencies in the forebrain, midbrain, hindbrain and neural tube but not in other tissues (from 1.3-fold in the neural tube (*P* = 7.3 × 10^-11^) to 1.6-fold in the forebrain (P = 1.03 × 10^-42^); Fig. 4a,c,d and Extended Data Fig. 6a,h,f). For example, the hs654 enhancer predominantly contacted *Zic1* and *Zic4* genes in the brain, neural tube and tail, tissues where enhancer and gene were both active (Figs. 3a and 4b). Interaction between hs654 and *Zic1/Zic4* was largely absent in face, limbs and heart tissues where both hs654 and *Zic1/Zic4* are inactive (Fig. 4b)^35^. Similarly, limb-specific enhancers displayed higher interaction frequencies with promoters in limb tissue (1.62-fold, *P* < 1.5 × 10^-37^), heart-specific in the heart (1.3-fold, *P* = 4.3 × 10^-09^), and face-specific in the face (1.62-fold, *P* = 3.6 × 10^-27^) (Fig. 4a,d and Extended Data Fig. 6a). We observed this pattern – that enhancers form significantly more frequent interactions with their respective target promoters when enhancers are active – for most enhancers in eight out of ten examined tissues (Fig. 4d and Extended Data Fig. 6a). There was no significant difference in interaction frequency for enhancers active in the tail and trunk, likely due to the low number of enhancers with characterized activity in these tissues (Extended Data Fig. 6a). We observed no significant increase in enhancer interactions with negative control regions in tissues where enhancers are active confirming the specificity of observed E—P interactions (Extended Data Fig. 6e).

We observed a similar trend even within developmentally related tissues, such as different parts of the brain. Enhancers active only in specific areas of the developing brain, formed significantly more frequent interactions with promoters in those tissues compared with parts of the brain where those enhancers were inactive (1.68-fold in the forebrain (*P* = 3.5 × 10^-8^) and 1.19-fold in the hindbrain (*P* = 0.027)) with the exception of the midbrain (Extended Data Fig. 6i,j). Notably, a small fraction of enhancers that formed invariant interactions with promoters across all tissues displayed an increased frequency of these interactions in tissues where the enhancer was active *in vivo* (Extended Data Fig. 7a,b). These results indicate that developmental gene activation is generally associated with an increased interaction frequency between corresponding enhancers and their target promoters.

We next examined *in vivo* chromatin interactions between enhancers (E—E contacts), including enhancers predicted based on chromatin features such as H3K27ac. Previous studies suggest a model in which enhancers regulating the same gene in the same cell form multi-enhancer hubs to activate gene expression^17,36,37^. We observed that E—E contacts formed between enhancers with overlapping activities are likely to regulate the same gene (Extended Data Fig. 3). For example, the hs268, hs267, hs266 and hs853 enhancers, which are located in the same TAD, formed extensive significant interactions with the promoter of the *mir9-2* gene (Extended Data Fig. 3a). All four enhancers were active in the dorsal telencephalon, and their activity patterns were strikingly similar to the expression of the *mir9-2* precursor (Extended Data Fig. 3a, c). All four enhancers also formed extensive interactions with each other in the forebrain (Extended Data Fig. 3a), but these E–E interactions were virtually absent in developing limb buds where *mir9-2* is not expressed, suggesting that these four enhancers form a multi-enhancer hub (Extended Data Fig. 3b). We observed similar tissue-specific E–E interactions at other loci and tissues (Extended Data Fig. 3d,e). Generally, enhancers formed significantly stronger interactions with other enhancers when they were active in the brain, face or limb (Fig. 4e,f and Extended Data Fig. 8b). These results are consistent with a model in which increased interactions among multiple enhancers during mammalian development and a given promoter accompanies transcriptional activation.

### Decrease in E—P distance in tissues where enhancers are active

To test whether the observed increase in E—P interactions also results in a change in a physical distance between enhancers and promoters^38,39^, we used super-resolution microscopy in conjunction with fluorescence in situ hybridization on three-dimensionally preserved nuclei (3D-FISH) to visualize enhancers and promoters in the developing mouse embryos. We chose three independent genetic loci where enhancer capture Hi-C revealed tissue-specific interactions between enhancers and their target genes (*Zic1/Zic4*, Fig. 4b; *mir9-2*, Extended Data Fig. 3a; *Snai2*, Fig. 6a). For all three genetic loci, the regulatory connection between enhancers and corresponding target genes was independently confirmed using enhancer knockout experiments (Fig. 3)^29^.

We performed 3D-FISH in forebrain, midbrain, craniofacial mesenchyme and forelimb cells at embryonic day E11.5 using fosmid-based probes targeting hs654, hs266 and hs1431 enhancers and corresponding target promoters. We observed a significant decrease in inter-probe distance (*P* = 1.18 × 10^-4^, hs654-*Zic1/Zic4* pair; *P* = 9.53 × 10^-7^, hs266-*mir9-2* pair; *P* =0.0106, hs1431-*Snai2* pair) and an increase in the fraction of co-localized alleles in tissues where corresponding enhancers are active for all three genetic loci (Fig. 5a,b and Extended Data Fig. 6n-p). For example, for hs266-*mir9-2* pair, the fraction of alleles with inter-probe distances less than 250 nm was 20% in the forelimb and increased to 32% in the forebrain (*P* = 1.47 × 10^-3^) where *mir9-2* is active (Fig. 5b and Extended Data Fig. 6o). A similar trend was observed for hs654-*Zic1/Zic4* pair (28% in the midbrain vs. 20% in the forelimb; *P* = 0.0132) and for hs1431-*Snai2* pair (32% in the face vs. 24% in the forebrain; not significant) (Fig. 5b and Extended Data Fig. 6n,p). Taken together, our 3D-FISH experiments showed a significant decrease in E—P physical distance in tissues where enhancers are active, which supports the increase in E—P interactions observed in our proximity-ligation-based enhancer capture Hi-C experiments.

**Fig. 5:**
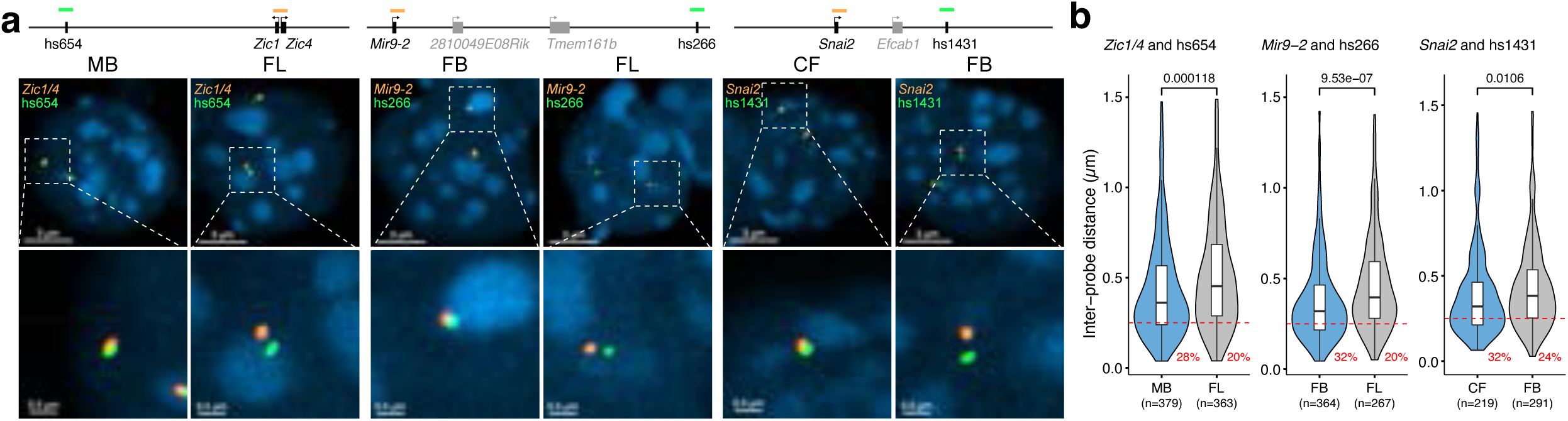
Imaging enhancer—promoter interactions in developing mouse embryo. **a**, The genomic positions of probes labeling enhancers (green) and genes (orange) are shown on the top. Gene and enhancer models are not drawn to scale. Images of representative nuclei (DAPI, blue) from E11.5 midbrain (left) and forelimb (right) after FISH with *Zic1*/4 and hs654 probe pairs (left panel), E11.5 forebrain (left) and forelimb (right) after FISH with Mir9-2 and hs266 probe pairs (middle panel), E11.5 face (left) and forebrain (right) after FISH with *Snai2* and hs1431 probe pairs (right panel) are shown. Corresponding zoomed in images are shown below. **b**, Violin plot showing the distribution of inter-probe distance (µm) between fosmid probe pairs in active and inactive tissues. Red dashed line indicates co-localization (<0.25 µm) and the numbers below represent the fraction of loci with co-localized probes. *P* values were calculated by paired-sample two-sided Wilcox test and adjusted for multiple testing for interaction frequencies comparison between active and inactive tissues, unpaired-sample two-sided Wilcox test was performed on comparison of inter-probe distance between different tissues. FB, forebrain. MB, midbrain. CF, craniofacial mesenchyme. FL, forelimb. For the boxplots in panel **b**, the central horizontal lines are the median, with the boxes extending from the 25th to the 75th percentiles. The whiskers further extend by ±1.5 times the interquartile range from the limits of each box.

### Properties of Invariant E–P interactions

Widespread stable mammalian E–P loops have been reported for enhancers, predicted from chromatin features in mouse embryonic limb and brain^21^, mouse embryonic stem cells^40,41^, and human keratinocytes^18^. However, how common is stable E–P looping at most developmental loci is unknown. Our analysis of E–P chromatin interactions for *bona fide* developmental enhancers found that only a small fraction (13.3%) formed tissue-invariant loops across all ten examined embryonic tissues (Fig. 6a-d). Nevertheless, these invariant E–P interactions displayed higher interaction frequency in tissues where enhancers were active (Extended Data Fig. 7a,b), consistent with increased E—P colocalization in transcriptionally active cells observed at preformed *Shh*/ZRS locus^42^.

**Fig. 6:**
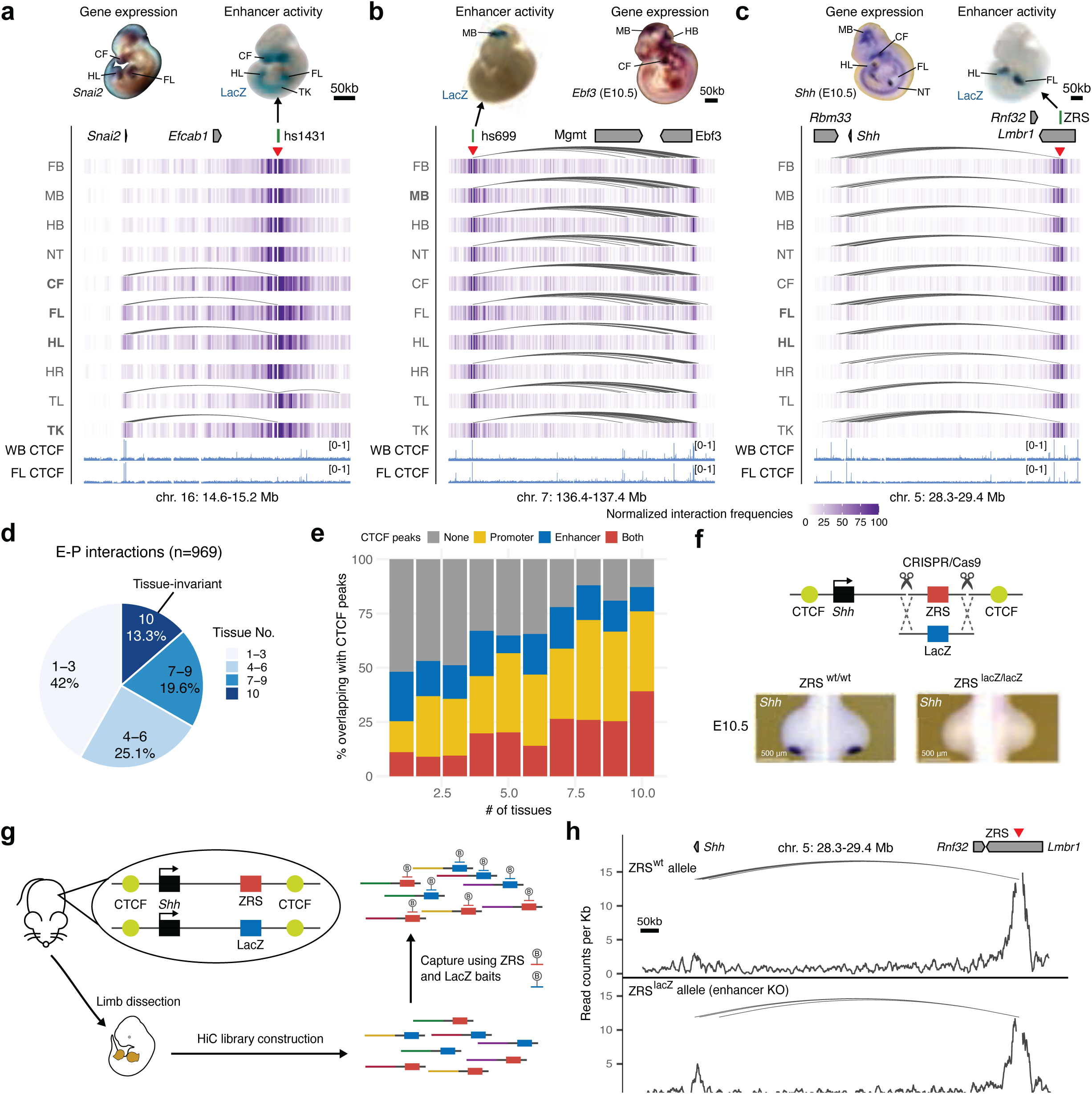
Properties of tissue-invariant enhancer–promoter chromatin interactions. **a-c**, Chromatin interaction profiles across 10 tissues centered on the hs1431 enhancer in the *Snai2* locus (chr16:14,610,000-15,220,000; mm10) (**a**), the hs699 enhancer in the *Dlx5/Dlx6* locus (chr7: 136,400,000-137,400,000; mm10) (**b**) and the ZRS enhancer in the *Shh* locus (chr5:28,320,000-29,400,000; mm10) (**c**). Shown above are corresponding enhancer activities in transgenic E11.5 mouse embryos and corresponding interacting gene mRNA WISH in E11.5 or E10.5 embryos. Heatmaps with normalized interaction frequencies in each of the 10 tissues are shown below. CTCF ChIP-seq profiles (blue) in the whole brain (WB) and forelimb (FL) at E12.5 are shown at the bottom^77^. Arches indicate significant interactions. Red arrowheads depict capture Hi-C viewpoints. **d**, Pie chart showing the fraction of E–P interactions present in different numbers of tissues. **e**, Fraction of E–P interactions that overlap with CTCF peaks grouped by number of tissues in which interaction was detected. f, Schematic of the Cas9-mediated strategy for replacement of the mouse ZRS sequence (red box) with a fragment of bacterial LacZ gene (blue box) at the *Shh* (black) genomic locus. CTCF binding sites are indicated in yellow. *Shh* mRNA WISH analysis in wild type and ZRS^LacZ/LacZ^ E10.5 mouse forelimb buds are shown below. See Extended Data Fig. 9 for details. g, Schematic overview of the capture Hi-C approach to detect chromatin interactions in the presence and absence of the ZRS in limbs of the same mouse using biotinylated RNA probes (B) targeting ZRS and LacZ. Limb buds from heterozygous transgenic mice were dissected followed by Capture Hi-C to enrich for ZRS and LacZ interactions. **h**, Allele-specific ZRS-region-centric chromatin interactions in limb buds of E11.5 ZRS^+/LacZ^ mice. Arches indicate significant interactions. WISH images in A and B have been reproduced with permission from Gene Expression Database (GXD, *Ebf3*)^35^ and Embrys database (http://embrys.jp; *Snai2*).

Stable E–P chromatin interactions are typically associated with neighboring CTCF binding^21,41^, especially for long-range E–P contacts such as ZRS-*Shh*^40,43^. Indeed, we observed that tissue invariant interactions are also associated with proximal CTCF binding, with more than 85% of all invariant interactions having proximal (< 5 kb) CTCF binding at either end, including the ZRS-*Shh* locus (Fig. 6c,e). By comparison, less than < 50% of tissue-specific interactions overlapped CTCF (Fig. 6e). The vast majority (87 out of 98, 88.8%) of enhancers that formed invariant interactions were active only in a subset of tissues similar to enhancers that form tissue-specific contacts (Extended Data Fig. 7c) which is consistent with a model in which CTCF forms these invariant interactions independently of enhancer activity.

To test if tissue-invariant interactions form independently of enhancer activity, we experimentally assessed how these E–P chromatin contacts are affected by targeted deletion of the enhancer. We chose the *Shh* locus where a limb-specific ZRS enhancer forms chromatin interactions with the *Shh* promoter located ∼850 kb away in all ten examined tissues (Fig. 6c). We generated a knock-in mouse line in which the entire ZRS enhancer was replaced with a piece of non-mouse DNA lacking any regulatory activity to simultaneously get rid of the enhancer and enable allele-specific detection of chromatin interactions in the capture Hi-C experiments. For that purpose, we used part of the bacterial *lacZ* gene sequence. Mice homozygous for the ZRS^lacZ^ allele showed no detectable *Shh* expression in the limb buds and displayed reduced limb buds at E11.5 and truncated zeugopods and autopods at E18.5, which is consistent with complete loss of *Shh* in the limb (Fig. 6f and Extended Data Fig. 9)^44^. To determine whether ZRS enhancer activity contributes to its higher-order chromatin interactions with the *Shh* promoter we performed capture Hi-C experiments in fully developed limb buds of E11.5 mice heterozygous for the ZRS^lacZ^ allele. Using probes targeting both the wild-type ZRS and LacZ sequence, we found that both the wild-type ZRS allele and “enhancerless” lacZ allele formed significant interactions with the *Shh* promoter (Fig. 6h). These results demonstrate that the higher-order chromatin interaction between ZRS and *Shh* can form independently of ZRS enhancer activity.

## Discussion

In this study, we comprehensively determined the tissue-resolved *in vivo* interaction landscapes for 935 *bona fide* enhancers, thus identifying thousands of tissue-specific interactions. Enhancer 3D chromatin conformations are highly dynamic across tissues and mirror the highly tissue-specific activity patterns observed for these enhancers in transgenic mouse embryos. We find moderate but consistent increases in E–P and E–E interactions in tissues where enhancers are functionally active. Together, our chromatin interaction data for 935 enhancers suggest that E–P physical proximity is a general feature of developmental gene activation in mammals.

Notably, we also detected E–P chromatin interactions that are tissue-invariant and are associated with proximal CTCF binding. Similar stable loops have been reported for other mammalian loci^18,21,22,43^ where it likely provides an additional level of robustness to maintain stable levels of gene expression during development^43^. Our data on *bona fide* enhancers suggests that these interactions occur next to a smaller fraction of developmental enhancers and likely form independently of enhancer activity. Since both tissue-invariant CTCF/cohesin-bound loops formed by loop extrusion and enhancer loops are widespread in the genome^45^, it is plausible that many of them overlap. Indeed, we did not observe differences in tissue specificity, evolutionary DNA conservation, or classes of target genes between enhancers that form tissue-invariant chromatin contacts and enhancers that form tissue-specific chromatin interactions with their promoters (Extended Data Fig. 7c-e).

While an increase in E—P interactions is linked to gene activation, the average observed increase in E–P contact frequency between active and inactive tissues appears to be less than 1.5-fold (Fig. 4c), even though average changes in associated tissue-specific gene expression are ∼11-fold (Extended Data Fig. 6g). Several models have been proposed to explain this nonlinear relationship between E–P contact probability and transcription, including bistability, hysteresis, and transient two-state E–P interactions^46,47^. The association between direct E—P contact and transcription at the macromolecular level remains elusive as some genetic loci show no or reverse association between E—P physical distance and transcription^24–26^. At least some differences could be due to the different approaches used to measure E—P interactions. Hi-C-based methods are based on proximity ligation and can be biased by crosslinking efficiency, while imaging-based methods, such as FISH, measure E—P distance directly. The two approaches sometimes result in contradicting results^26,38,39,48^. Higher resolution imaging techniques and C-methods as well as methods based on live imaging will be needed to untangle complex relationships between direct E—P contacts and transcription^49–52^.

Our results contrast with other systems such as early Drosophila embryo development^20,53,54^ or stimulus-induced gene activation^55,56^ where E–P loops appear to be stable and are often associated with paused Pol II^20^. In these specialized systems, pre-formed E–P topologies might ensure robust and rapid gene activation^13,20^. Interestingly, the emergence of new E–P loops correlates with enhancer activation in differentiated *Drosophila* embryonic tissues, suggesting that E–P proximity could be an evolutionary conserved property of mid-late animal embryogenesis^57^.

More than half of developmental enhancers in our study appear to skip neighboring genes to regulate a more distal one. Such interactions have also been reported in mice^58,59^, human^60,61^, and to a lesser degree in *Drosophila*^62,63^. This raises the question: How is this E–P selectivity achieved? Our analysis of remote E–P interactions shows that promoters of approximately half of the skipped genes are methylated and inaccessible (Fig. 2d-f and Extended Data Fig. 4), suggesting that promoter silencing could potentially be one of the mechanisms by which such enhancer–gene specificity is achieved in mammals^64^. However, the other half of promoters skipped by distal enhancers are not methylated and are accessible at comparable levels with target genes indicating that additional factors facilitate promoter bypassing by remote enhancers. Such factors could potentially include compatibility between enhancers and different types of core promoters^65–68^ and tethering elements^63,69,70^. The general mechanism that determines E–P specificity in mammalian genomes is still poorly understood^71^, and further studies are needed to dissect how divergent expression is achieved within the same TAD. Notably, we also observe that 21% of developmental enhancers act across TAD boundaries confirming previous observations^72,73^. These cross-TAD enhancers behave similarly to intra-TAD enhancers (Extended Data Fig. 6c) but tend to locate closer to TAD borders (Extended Data Fig. 6d) consistent with the boundary staking model that was proposed to facilitate TAD border bypass^73^.

It is important to note that the current study surveyed a relatively small fraction of *bona fide* developmental enhancers in a limited number of mouse embryonic tissues and timepoints. In future studies, functional characterization of a greater number of developmental enhancers and their chromatin interactions *in vivo* in various tissue and cell contexts will greatly aid functional interpretation of germline variants associated with human congenital disorders. Nonetheless, the current study provides a broad snapshot of the general 3D chromatin organization and properties of enhancers at typical developmental loci.

## Acknowledgments

The authors would like to acknowledge the UCI Transgenic Mouse Facility for help with generation of enhancer knockout mice and the UCI Genomics Research and Technology Hub (GRT Hub) for help with sequencing, as well as Drs. Lorenzo Scipioni and Sha Sun for help with DNA FISH. Funding: This work was supported by National Institutes of Health grants R00HG009682 and DP2GM149555 (to E.Z.K.), R01HG003988 (to L.A.P.) and F31HD112201 (to G.B.). Z.C. was supported by NSF grant DMS1763272 (to Qing Nie) and Simons Foundation grant 594598 (to Qing Nie). J.L-R. is funded by the Spanish Ministerio de Ciencia e Innovación (grant PID2020-113497GB-I00 and institutional María de Maeztu grant CEX2020-001088-M). Research conducted at the E.O. Lawrence Berkeley National Laboratory was performed under Department of Energy Contract DE-AC02-05CH11231, University of California. The funders had no role in study design, data collection and analysis, decision to publish or preparation of the manuscript.

## Author contributions

E.Z.K. conceived the project with input from Z.C., V.S., D.E.D., A.V. and L.A.P. Z.C., V.S., I.B. and E.Z.K. designed experiments. Z.C., V.S., G.B., S.J., B.C. and E.Z.K. performed Capture Hi-C experiments and Z.C. analyzed the data with input from E.Z.K. and B.J.M. Z.C., S.J., A.D. and E.Z.K. performed the enhancer knockout studies and Z.C. analyzed the data. Z.C. and G.B. performed 3D-FISH experiments and analyzed the data. A.A.C. and J.L-R. performed ISH experiments. E.Z.K and Z.C wrote the manuscript with input from the remaining authors.

## Extended Figure Legends

**Extended Data Fig. 1.**
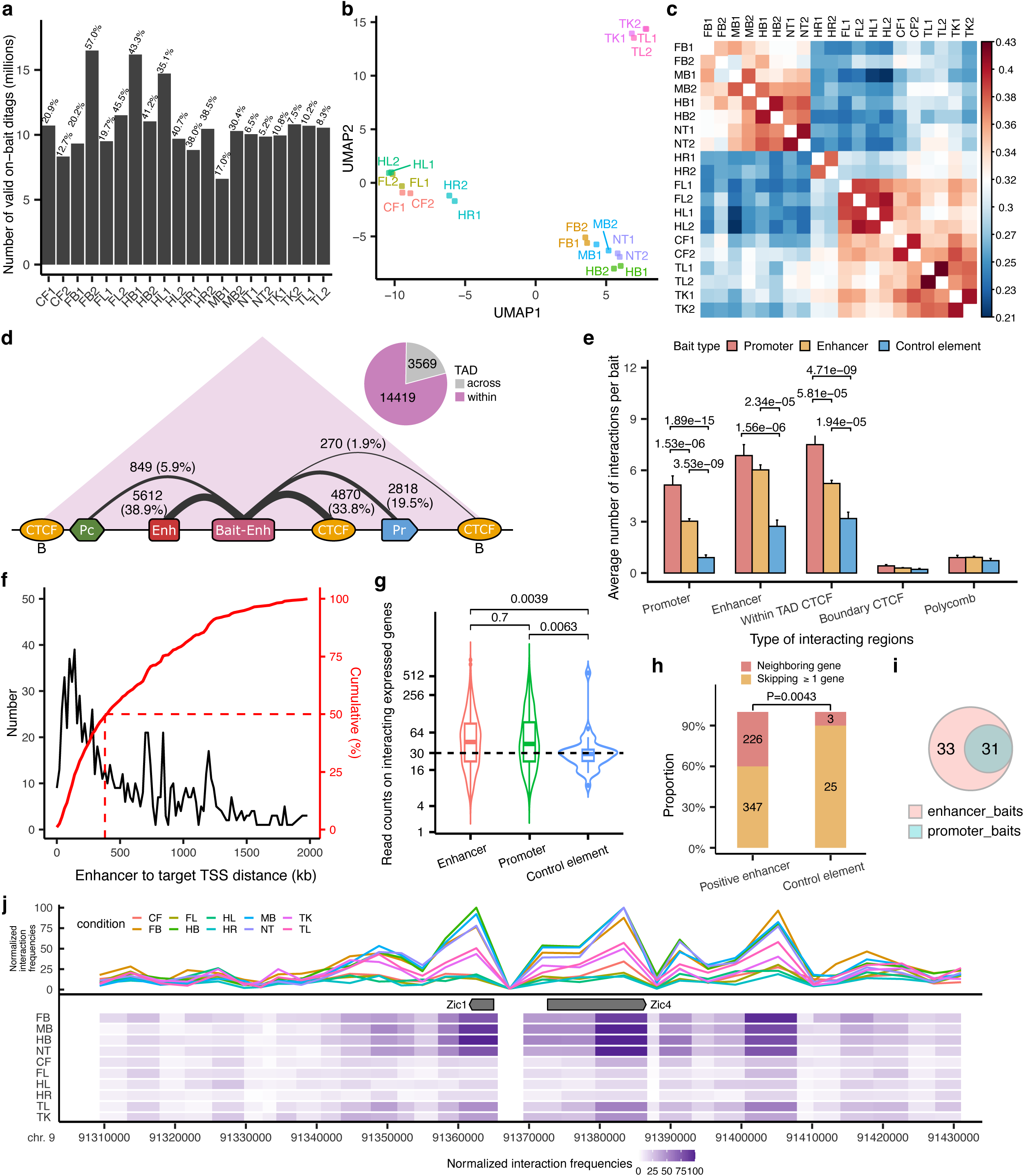
Enhancer capture Hi-C identifies enhancer-centric chromatin interactions in mouse embryonic tissues. **a**, Unique on-target read counts for each library. The percentages above indicate the capture rates for each library. **b,c**, Principal component analysis and hierarchical clustering of all replicates based on the presence of peaks called by CHiCAGO in each replicate (considering peaks with valid di-tags on neighboring fragments). **d**, Significant enhancer-centric chromatin interactions identified in this study. The number on each link represents the number of fragments falling into different annotation categories and the width of links is proportional to the percentage (in the parentheses) of different kinds of interactions. Only interactions within 2 Mb are included. CTCF sites with “B”: CTCF sites at TAD boundary; Pc: polycomb; Enh: enhancers; Bait-Enh: baited enhancers; Pr: promoters. **e**, An average number of interactions detected per bait for different kinds of baits (promoter (*n*=176), enhancer (*n*=935) and negative control elements (*n*=87)). Data are represented as mean ± s.e.m. **f**, Distribution of genomic distances between enhancers and the TSSs of interacting genes (black, frequencies; red, cumulative). **g**, Violin plots showing read counts on promoters of active genes that interact with enhancer baits (*n*=541), promoter baits (*n*=126) and control element baits (*n*=25). The central horizontal lines are the median, with the boxes extending from the 25th to the 75th percentiles. The whiskers further extend by ±1.5 times the interquartile range from the limits of each box. **h**, Histogram showing the proportion of bait regions that interact with proximal genes and distal genes. **i**, Venn diagram showing the overlap between significant interactions called from enhancer baits and corresponding promoter baits. All *P* values were calculated by a two-sided Wilcox test and adjusted for multiple testing. **j**, Zoom-in view on *Zic1/Zic4* locus for hs654 interaction profiles across 10 tissues. The average size for each pooled fragment is ∼3kb. FB, forebrain. MB, midbrain. HB, hindbrain. CF, craniofacial mesenchyme. HR, heart. FL, forelimb. HL, hindlimb. TK, trunk. TL, tail. NT, neural tube.

**Extended Data Fig. 2.**
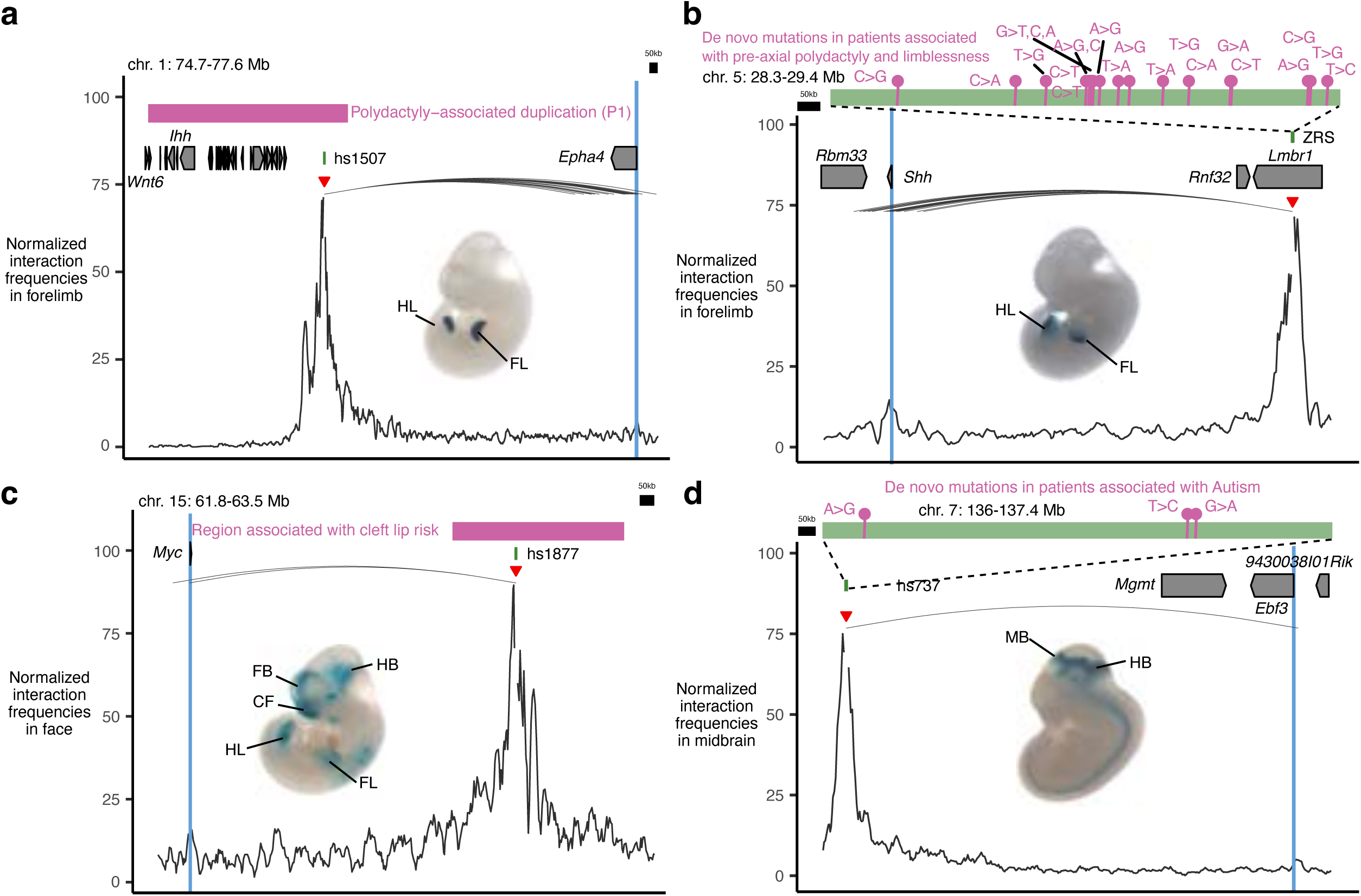
Examples of enhancer—promoter interactions linked to congenital disorders. **a**, Hs1507 limb enhancer located in the non-coding region which is duplicated in patients with polydactyly (pink box indicates the homologous region in the mouse genome)^8^. Hs1507 forms significant chromatin interactions with the promoter of the *Epha4* located ∼1.5 Mb away. Shown is the *Epha4* genomic region (chr1:74,788,119-77,634,678; mm10). **b**, Many *de novo* rare variants identified in patients with preaxial polydactyly^103^ are located in the ZRS limb enhancer which forms significant interactions with the promoter of *Shh* located ∼850 kb away. Shown is the *Shh* genomic region (chr5:28,320,000-29,400,000; mm10). **c**, Hs1877 face enhancer located in the non-coding region containing 146 SNPs found in patients with cleft lip risk^104^ (pink box indicates the homologous region in the mouse genome). Hs1877 forms significant chromatin interactions with the promoter of the *Myc* located ∼900 kb away in the face. The *Myc* genomic region (chr15:61,880,003-63,506,895; mm10). **d**, Three *de novo* rare variants identified in patients with autism are located in the hs737 midbrain/hindbrain enhancer^105,106^, which forms strong significant interactions with the promoter of *Ebf3* located ∼1,000 kb away in the midbrain. Shown is the *Ebf3* genomic region (chr7:136,018,204-137,420,338; mm10).

**Extended Data Fig. 3.**
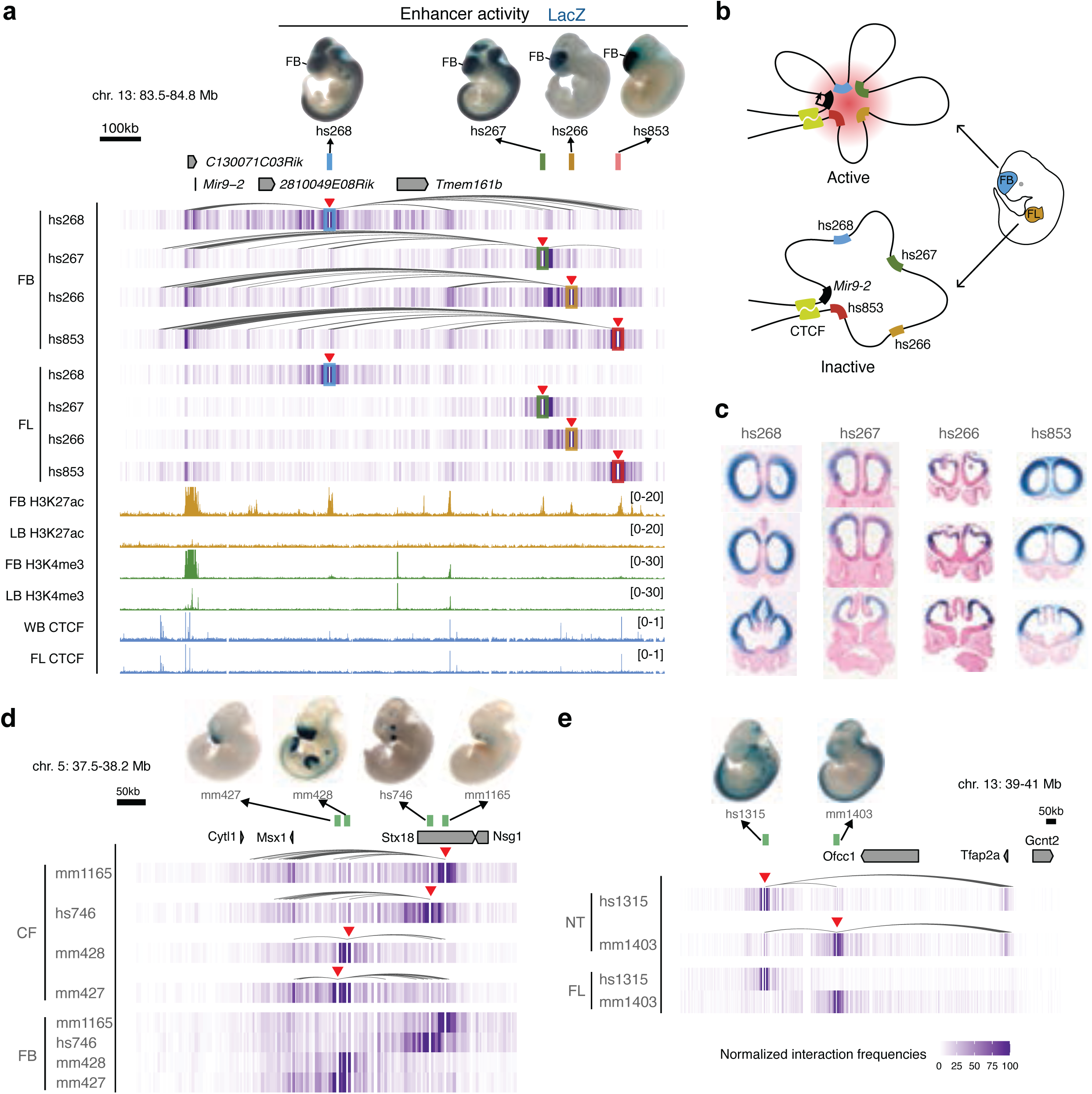
Examples of enhancer—enhancer chromatin interactions. **a**, The *Mir9-2* genomic region (chr13:83,558,457-84,861,438; mm10) is shown with chromatin interaction heatmaps centered on hs268 (blue), hs267 (green), hs266 (yellow) and hs853 (red) enhancers in the forebrain (FB) and forelimb (FL). Shown on the top are hs268, hs267, hs266 and hs853 enhancer activities in a transgenic mid-gestation (E11.5) mouse embryo, which match with the expression profiles of *Mir9* in the brain and neural tube at E11.5^107,108^. Red arrowheads indicate capture Hi-C viewpoints. Arches indicate significant interactions in the forebrain. Shown on the bottom are H3K27ac (yellow) and H3K4me3 (green) ChIP-seq tracks in forebrain and limb buds (LB) at E11.5, CTCF (light blue) ChIP-seq tracks in the whole brain (WB) and forelimb at E12.5^34,76,77,109^. **b**, Schematic depicting 3D chromatin interactions between enhancers and *Mir9-2* gene in the forebrain and forelimb. c, Coronal sections of forebrain for hs268, hs267, hs266 and hs853 enhancer activity from VISTA enhancer database^28^, which reproducibly label the same subregions in E11.5 forebrain as *C130071C03Rik* (*Mir9-2* precursor) expression^108^. **d,e**, Chromatin interaction heatmaps centered on mm1165, hs746, mm428 and mm427 enhancers in the face (CF) and forebrain (FB) for Msx1 genomic region (chr5: 37,554,764-38,206,723; mm10) (**d**) and hs1315 and mm1403 enhancers in the neural tube (NT) and forelimb (FL) for *Tfap2a* genomic region (chr13: 39,098,000-41,000,000; mm10) (**e**). Shown on the top are mm1165, hs746, mm428, mm427, hs1315 and mm1403 enhancer activities in a transgenic mid-gestation (E11.5) mouse embryos. Arches indicate significant interactions.

**Extended Data Fig. 4.**
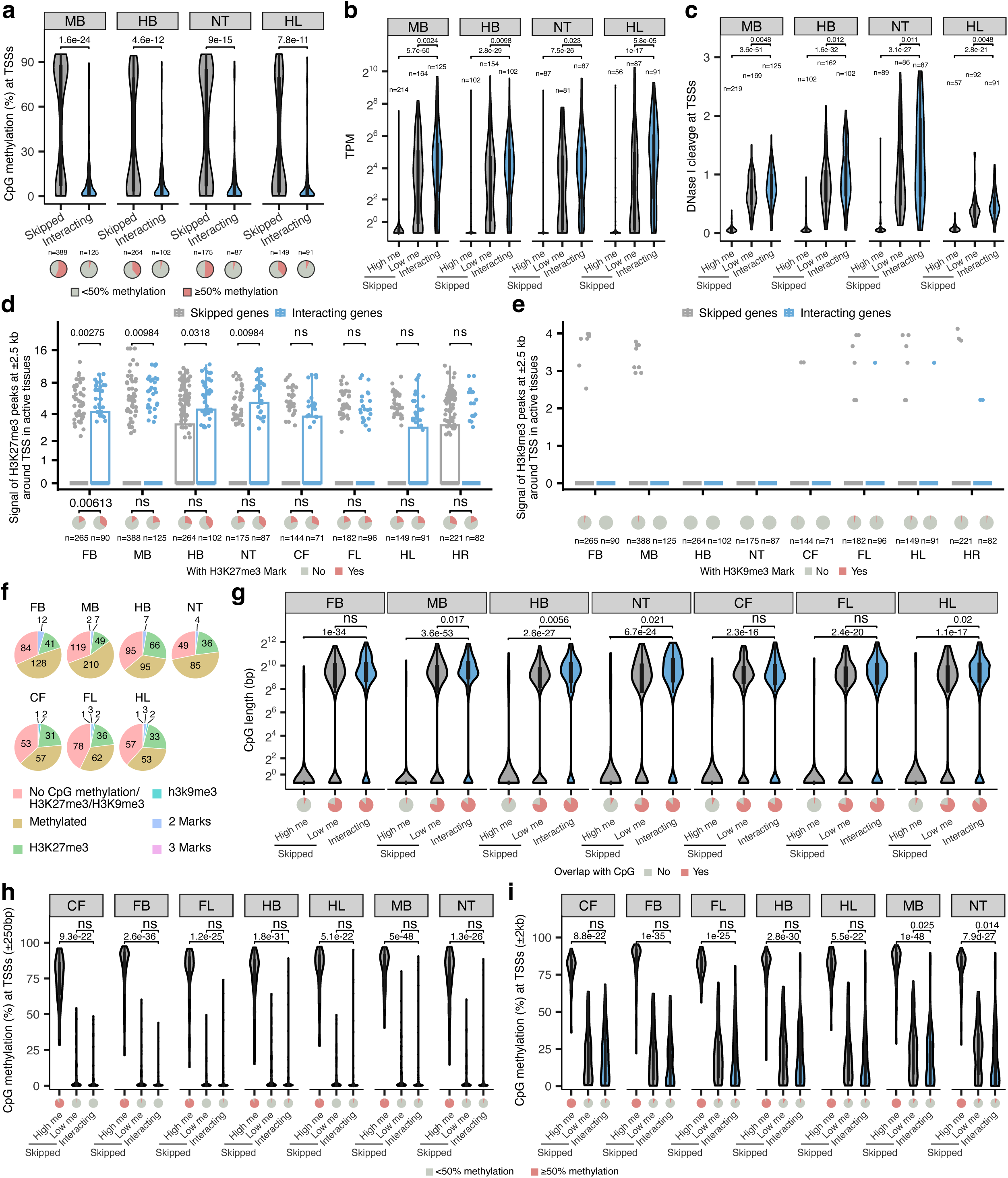
Properties of enhancer-interacting and skipped promoters. **a-c**, The CpG methylation (**a**), mRNA expression levels (**b**) and DNase signal (**c**) of enhancer-interacting and skipped promoters in tissues where enhancers are active. High me, high methylation skipped promoters (>50% CpG methylation within ± 1 kb from TSS). Low me, low methylation skipped promoters (<50% CpG methylation within ± 1 kb from TSS). **d,e**, H3K27me3 (**d**), H3K9me3 (**e**) signal at ± 2.5 kb of enhancer-interacting and skipped promoters in tissues where enhancers are active. The pie charts below show the fraction of promoters marked with H3K27me3 or H3K9me3. **f**, Pie charts showing the fraction of skipped promoters marked by CpG methylation, H3K27me3, H3K9me3 or the combination of marks. g-i, Violin plot showing CpG length (g), or CpG methylation level at transcription start sites for enhancer-interacting and skipped genes with different window sizes ± 250bp (**h**) and ± 2kb (**i**)). The number of high and low methylated skipped as well as interacting promoters in CpG analysis are *n* =58, *n* =86 and *n* =71 (CF), *n* =138, *n* =126 and *n* =90 (FB), *n* =64, *n* =116 and *n* =96 (FL) and *n* =100, *n* =162 and *n* =102 (HB), *n* =55, *n* =92 and *n* =91 (HL), *n* =213, *n* =169 and *n* =125 (MB) and, *n* =87, *n* =86 and *n* =87 (NT). FB, forebrain. MB, midbrain. HB, hindbrain. CF, craniofacial mesenchyme. FL, forelimb. HL, hindlimb. NT, neural tube. HR, heart. *P* values are calculated by two-sided Wilcoxon rank test after adjusted for multiple testing (a-c, f-i) or by one-sided chi-squared test (d, e). A statistical test was not performed for H3K9me3 since most of the values are zero. The same DNA methylation, mRNA expression, DNaseI hypersensitivity, H3K27ac and H3K9me3 dataset (a mixture of fore- and hindlimb buds) were used for both fore- and hindlimb interaction analyses. For the boxplots in panels a-e and g-i, the central horizontal lines are the median, with the boxes extending from the 25th to the 75th percentiles. The whiskers further extend by ±1.5 times the interquartile range from the limits of each box.

**Extended Data Fig. 5.**
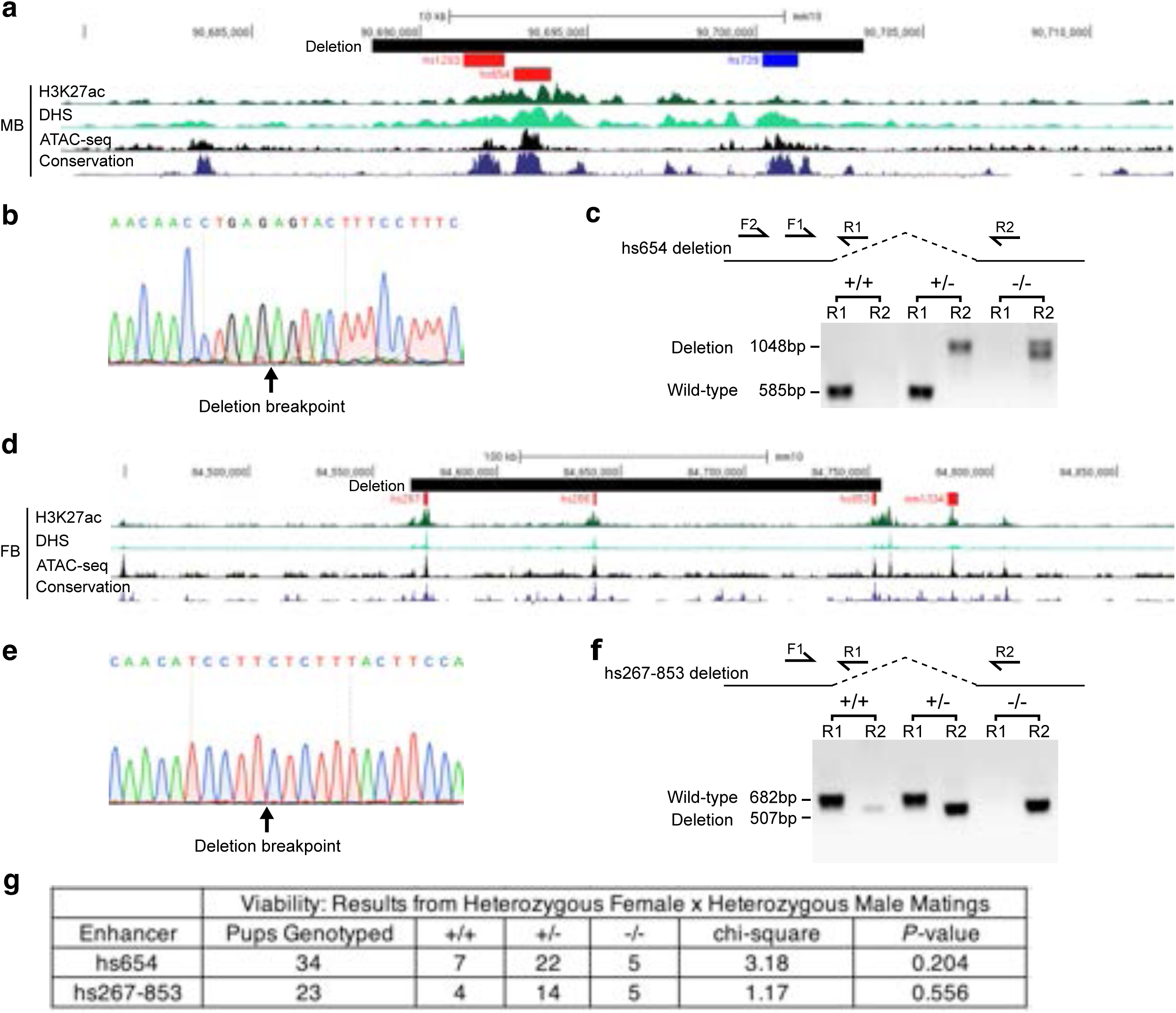
*Zic1/Zic4* and *Mir9-2* brain enhancer knock-outs. **a**, Map of the deleted region encompassing hs654 midbrain enhancer of *Zic1/Zic4* together with H3K27ac, DNase-seq, ATAC-seq from midbrain and conservation track across 60 species. **b**, Sanger sequencing of the PCR product from hs654 knock-out mice (*n* = 4 biological replicates). **c**, representative PCR genotyping results of the hs654 enhancer knockout mice. Lanes in the gel were rearranged so that results for wild-type and heterozygous mice are adjacent to each other. **d**, Map of the deleted region encompassing hs267, hs266 and hs853 forebrain enhancers of *Mir9-2* together with H3K27ac, DNase-seq, ATAC-seq from midbrain and conservation track across 60 species. **e**, Sanger sequencing of the PCR product from hs267-853 knock-out mice (*n* = 3 biological replicates). **f**, representative PCR genotyping results of the hs267-853 enhancer knockout mice. **g**, Genotype frequency data for enhancer knockout lines. Mice homozygous for either deletion were born at normal Mendelian ratios, and no gross phenotypes or impairments were observed. *P*-values were calculated using the one-sided chi-square test. **h**, Primer sequences used for genotyping of enhancer knock-out mice.

**Extended Data Fig. 6:**
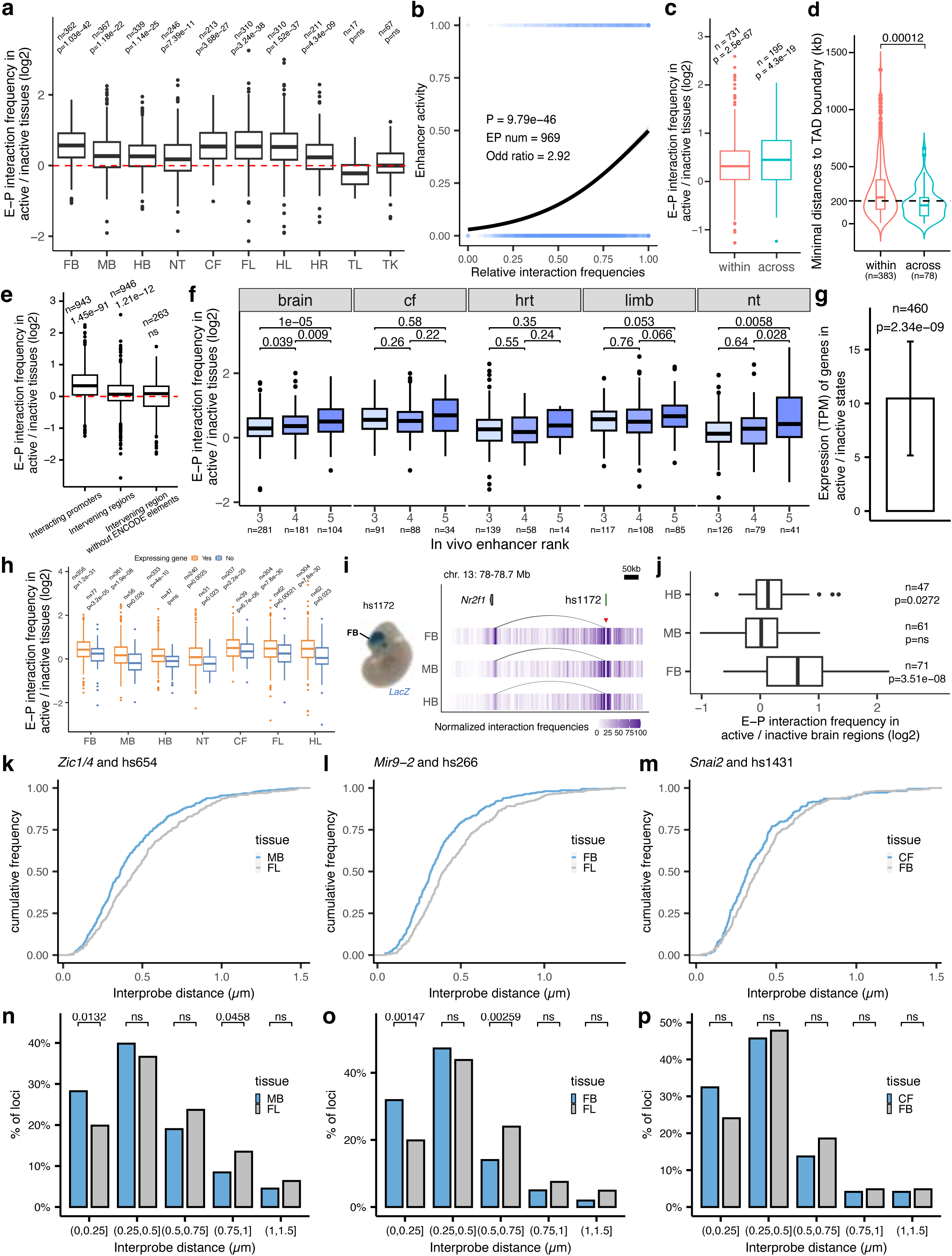
E–P interaction frequency in active and inactive tissues. **a**, The ratio of E–P interaction frequency between active and inactive tissues. **b**, Univariate logistic regression for relative interaction frequencies and enhancer activity across all tissues. **c**, The ratio of E–P interaction frequency between active and inactive tissues for interactions within or across TADs. **d**, The distribution of distances between the closest TAD boundary and enhancer for enhancers acting within or across TADs. **e**, The ratio of interaction frequency between active and inactive tissues on interacting promoters or intervening regions before and after removing ENCODE annotated elements (±20kb). **f**, The ratio of E–P interaction frequency between active and inactive tissues for enhancers with different ranks. Only tissues with ≥10 interactions in each rank category are shown. **g**, The fold-change of gene expression levels between active state (baited enhancers interact with active promoters) and inactive state (baited enhancers don’t interact with promoters or in inactive tissues). Data are represented as mean ± s.e.m. **h**, The ratio of E–P interaction frequency between active and inactive tissues for expressed genes (TPM>=0.5) and lowly expressed or inactive genes (TPM<0.5). i, Chromatin interaction profiles in forebrain, midbrain and hindbrain centered on the enhancer hs1172 at *Nr2f1* locus (chr13:78,057,768-78,705,499). **j**, The ratio of E–P interaction frequency between active and inactive brain regions for enhancers active in one of the brain domains. **k-m**, Cumulative frequency plots of inter-probe distances for the indicated loci and tissues. **n-p**, Frequency distribution of FISH inter-probe distances in 250 nm bins between *Zic1/4* and *hs654* (**n**), *Mir9-2* and hs266 (**o**), Snai2 and hs1431 (**p**) in indicated tissues. *P* values are calculated by paired-sample (a, c, e, g, h, j) or unpaired-sample (d, f) two-sided Wilcoxon rank test and adjusted for multiple testing or by one-sided chi-squared test (b, n-p). For the boxplots in panels a, c-f, h and j, the central horizontal lines are the median, with the boxes extending from the 25th to the 75th percentiles. The whiskers further extend by ±1.5 times the interquartile range from the limits of each box.

**Extended Data Fig. 7:**
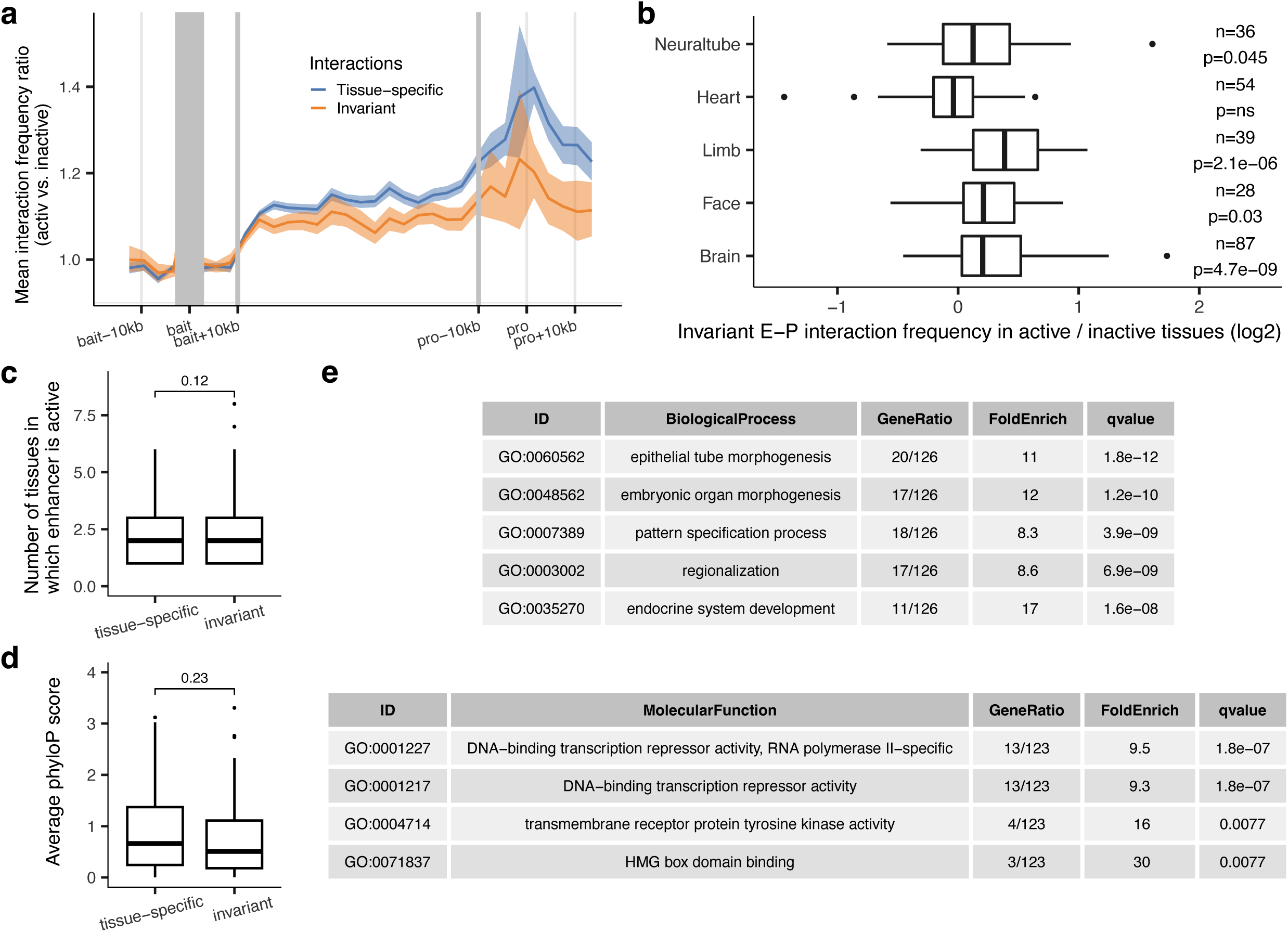
Properties of invariant E–P interactions. **a**, Metaplot showing average ratio of enhancer interaction frequency between active and inactive tissues for invariant (interactions present in all 7 main tissues: brain, face, limb, heart, neural tube, trunk and tail, *n*=171) and tissue-specific (≤ 6 main tissues, *n*=775) interactions. Light blue/orange shading indicates 95% confidence intervals estimated by non-parametric bootstrapping. **b**, The average ratio of invariant enhancer-promoter interaction frequency between active and inactive tissues for enhancers active in the brain, face, limb, heart and neural tube E–P. Data is shown only for tissues with at least 20 active enhancers that form invariant E–P interactions. *P* values were calculated by paired-sample two-sided Wilcox test and adjusted for multiple testing. **c**, The number of tissues in which enhancers forming invariant (10 tissues, *n*=98) or tissue-specific (≤ 4 tissues, *n*=196) E–P interactions are active *in vivo*. **d**, The average phyloP scores of enhancers forming invariant (10 tissues, *n*=98) or tissue-specific (≤ 4 tissues, *n*=196) E–P interactions. *P* values in panels c and d were calculated by two-sided Wilcox test. **e**, Gene Ontology enrichment for genes that form invariant (10 tissues) E–P interactions (Biological process and Molecular function). Q values were calculated by over-representation test and adjusted for multiple testing. For the boxplots in panels b-d, the central horizontal lines are the median, with the boxes extending from the 25th to the 75th percentiles. The whiskers further extend by ±1.5 times the interquartile range from the limits of each box.

**Extended Data Fig. 8:**
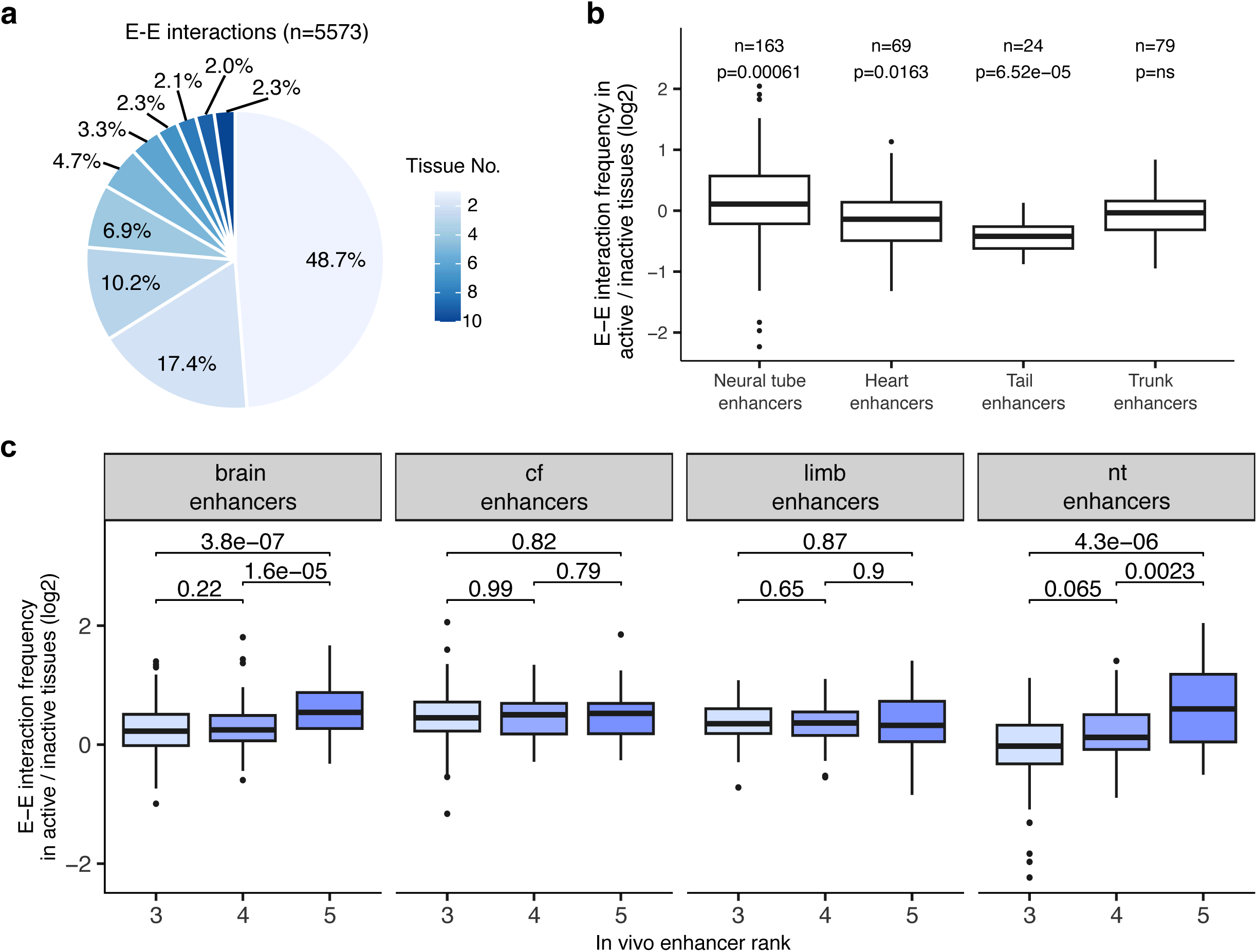
Tissue specificity of enhancer-enhancer chromatin interactions. **a**, Pie chart showing the fraction of E–E interactions present in different numbers of tissues. **b**, The average ratio of E–E interaction frequency between active and inactive tissues for enhancers active in neural tube, heart, tail and trunk. The number of E–E interactions for each tissue is indicated at the top. *P* values were calculated by paired-sample two-sided Wilcox test and adjusted for multiple testing. **c**, The average ratio of enhancer–enhancer interaction frequency between active and inactive tissues for enhancers of different ranks. The E–E interaction number for rank 3 to 5 are *n*=217, *n*=122 and *n*=69 (brain), *n*=53, *n*=59 and *n*=18 (cf), *n*=100, *n*=84 and *n*=45 (limb), *n*=80, *n*=51 and *n*=32 (nt), respectively. Cf: face. Nt: neural tube. *P* values were calculated by unpaired-sample two-sided Wilcox test with multiple testing. For the boxplots in panels b and c, the central horizontal lines are the median, with the boxes extending from the 25th to the 75th percentiles. The whiskers further extend by ±1.5 times the interquartile range from the limits of each box.

**Extended Data Fig. 9:**
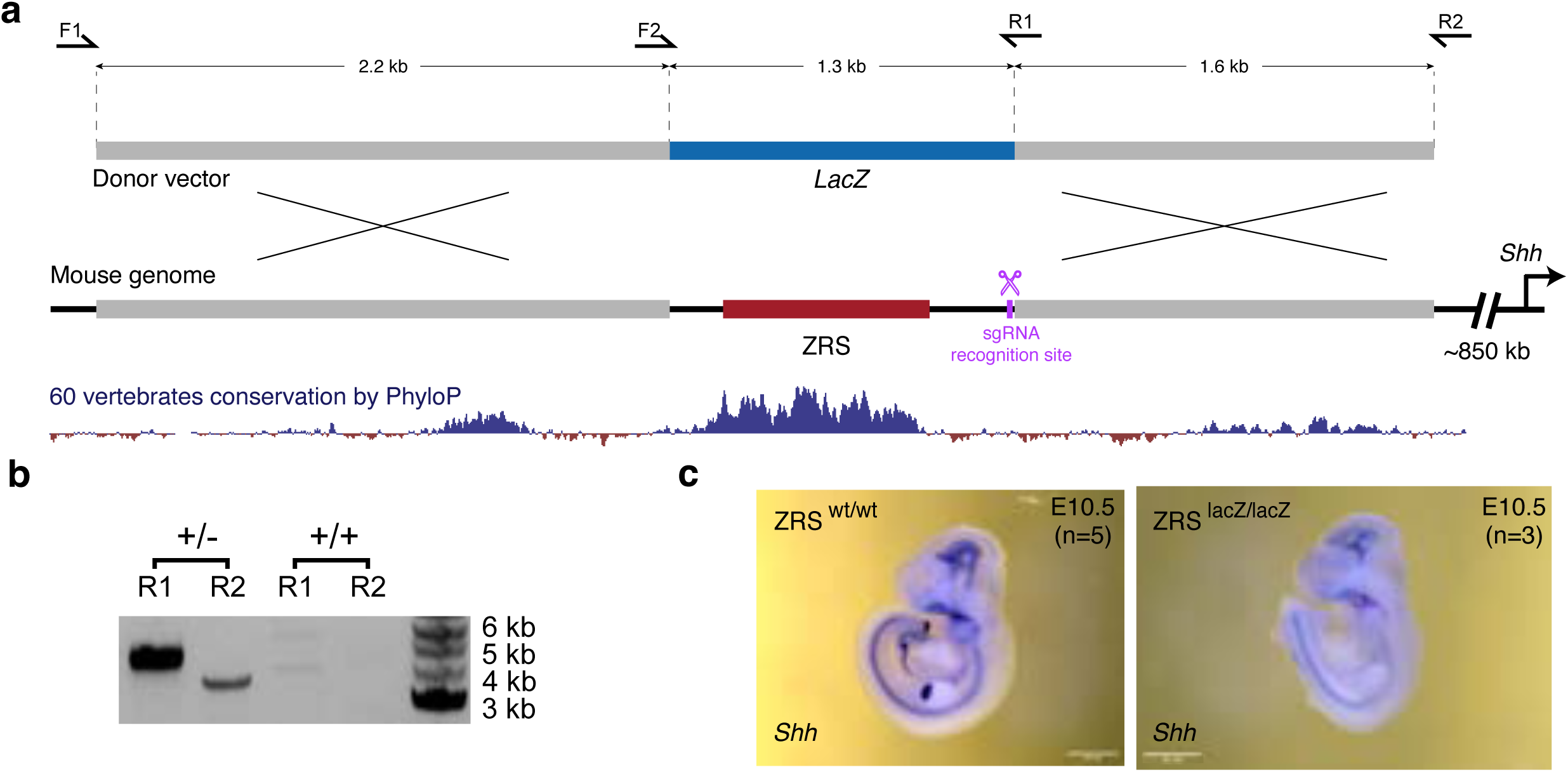
CRISPR/Cas9-mediated ZRS limb enhancer replacement with a fragment of the *lacZ* gene. **a**, Schematic overview of the strategy for ZRS enhancer replacement. A 4.5 kb mouse genomic region containing the ZRS enhancer (red) is shown together with the vertebrate conservation track (dark blue). The donor vector contained two homology arms (gray) and an inactive fragment of the *lacZ* coding sequence (blue). The sgRNA recognition site is indicated in purple. PCR primers used for genotyping are shown as arrows. **b**, PCR genotyping analysis of heterozygous and wildtype mice using primer pairs LacZ-F1 and LacZ-R1 or LacZ-F2 and LacZ-R2. See **Methods** for details. **c**, *Shh* whole-mount in situ hybridization in E10.5 wild type (left) and ZRS^lacZ/lacZ^ knock-in embryos (n ≥ 3 biological replicates for each genotype). *Shh* expression is not detectable in limb buds but is present elsewhere in the embryo. d, Primer sequences used for genotyping of ZRS^lacZ/+^ knock-in mice.

## Methods

### Ethics statement

All animal work was reviewed and approved by the Lawrence Berkeley National Laboratory Animal Welfare and Research Committee and the University California Irvine Laboratory Animal Resources (ULAR) under protocols AUP-20-001 and AUP-23-005. Mice were housed in the animal facility, where their conditions were electronically monitored 24/7 with daily visual checks by technicians.

### Tissue collection

Mouse embryonic tissues, including the forebrain, midbrain, hindbrain, neural tube, tail, facial mesenchyme, forelimb, hindlimb, heart and trunk, were collected from FVB/NCrl strain *Mus musculus* animals (Charles River). Wild-type male and female mice were mated using a standard timed breeding strategy and E11.5 embryos were collected for dissection using approved institutional protocols. Embryos were excluded if they were not at the expected developmental stage. Only one embryonic litter was processed at a time and tissues and embryos were kept on ice to avoid degradation during tissue collection. Tissue from multiple embryos was pooled together in the same collection tube, and at least two separate tubes were collected for each tissue for biological replication.

### Tissue processing for Hi-C library

To prepare nuclei for constructing the Hi-C library, tissues were incubated with collagenase (Gibco) in a thermomixer at 37°C until the cells were dissociated, about 10 to 20 min. Cells were fixed by adding formaldehyde (Sigma-Aldrich) to a final concentration of 2% at RT for 10 min^43,82^. Ice-cold glycine solution was added to a final concentration of 200 mM to quench crosslinking. Cells were then resuspended in cold lysis buffer (50 mM Tris, pH 7.5, 150 mM NaCl, 5 mM EDTA, 0.5% NP-40, 1.15% Triton X-100 and 1X protease inhibitor cocktail (Thermo Scientific)) and incubated on ice for 15 min. Pellets of nuclei were obtained by centrifuge at 750 g for 5 min at 4°C, followed by snap-freezing and storage at -80°C.

### Generation of Hi-C library

Hi-C libraries were prepared as described previously^82–84^. Briefly, frozen nuclei pellets (2-6 million) were thawed on ice, followed by adding SDS and Triton X-100 to remove non-crosslinked proteins and sequester SDS, and digested using DpnII (NEB) overnight at 37°C. The ends of restriction fragments were labeled with biotinylated dCTP and ligated at room temperature for 4 hours. After de-crosslinking and precipitation, ligated products were sheared using a Covaris sonicator (duty cycle: 10%, intensity: 5, cycles per burst: 200, treatment time: 180 s in total) to an average fragment size of 200bp. The ligated sheared 3C libraries (10-12 µg for each replicate) were pulled down using Streptavidin Dynabeads (Thermo Scientific) to get rid of unligated fragments, followed by end repair, adaptor ligation and library amplification according to modified Agilent SureSelectXT protocol.

### Capture Hi-C probe design

To perform enhancer capture Hi-C, we designed 120-mer RNA probes, targeting 935 enhancer regions that showed highly reproducible activity at E11.5 from VISTA Enhancer Database^85^ (**Supplementary Table 1**). We also designed RNA probes targeting 176 promoters and 87 elements with no reproducible enhancer activity at E11.5 as negative controls (**Supplementary Table 1**). All elements shorter than 2 kb were re-sized to 2 kb (± 1 kb from their central coordinate).

We designed 20,452 120-mer probes (each region was covered by on average 17 RNA probes) using the following pipeline. We first identified the DpnII restriction sites (GATC) overlapping each element by generating a genome-wide map of cut sites using vmatch (http://www.vmatch.de/). For each of the DpnII restriction sites overlapping the re-sized VISTA elements, ± 240 bp around the recognition site were considered for tiling. Among the resulting regions, those found within 60 bp of each other were further merged. After that, these regions were tiled (from -60 bp to +60 bp) using overlapping 120 bp windows, with a step of 60 bp. The tiles obtained were further filtered based on their overlap with repetitive elements and their predicted mappability using short reads. For filtering based on mappability, the wgEncodeCrgMapabilityAlign36mer.bigWig track from the UCSC genome browser (mm9) was used. Only tiles showing a mappability score of 1 across all 120 bp were retained. For exclusion based on repeats, the tiles were first lifted to mm10 (using liftOver), then each tile showing an overlap of at least 10% with an annotated repeat in the RepeatMasker track of the UCSC genome browser were excluded. Following that, only those overlapping elements represented by at least three tiles were considered for the final design. For capture Hi-C experiments at the *Shh*-ZRS locus (Fig. 6) we designed a separate panel that covered the ZRS enhancer, part of the bacterial LacZ sequence and 9 control regions (**Supplementary Table 1**).

### Capture Hi-C library construction and sequencing

The enhancer capture Hi-C library was created by performing a target-enrichment protocol using capture RNA probes according to Agilent SureSelect XT protocol with an input amount of 750 ng of Hi-C library per sample. Following hybridization to the RNA oligo library, each capture Hi-C library was sequenced (paired-end 100 or 150 bp) to enrich enhancer-centric interactions yielding a total of 1 billion unique paired-end reads.

### Capture Hi-C data analysis

After checking read quality by FastQC (v0.11.9), ligated reads were trimmed using DpnII restriction recognition sites and mapped to the DpnII-digested reference genome (mm10) using HiCUP (v0.8.0)^86^, followed by quality filtering and deduplication. For each tissue, the capture Hi-C experiment produced, on average, 20 million unique on-target paired-end reads, resulting in a total of 200 million valid read pairs (**Supplementary Table 1**).

Next, all DpnII fragments overlapping with the same bait region were merged into a single fragment *in silico*. Subsequently, the rest of the DpnII fragments were merged based on the size distribution of the pooled fragments that overlapped with bait regions. The mean fragment size of pooled fragments is ∼3,000 bp. Significant interactions were called by CHiCAGO (v1.26.0, score > 5) with the default setting^87,88^ using combined replicates from HiCUP pipeline, by using the design file with the following parameters: --minFragLe*n*=300 --maxFragLe*n*=20000 --binsize=20000 -- maxLBrownEst=3000000 --removeAdjacent=FALSE. We removed significant interactions that didn’t have valid di-tag reads on neighboring fragments to avoid spurious interaction spikes^89^. Interactions called >2 Mb from the bait regions were excluded from the downstream analysis.

To visualize and compare interaction frequencies between different tissues, read counts were normalized across 10 tissues by Chicdiff (v0.6)^88,90^ to account for library size and background differences between samples. We used the output from CHiCAGO to make a peak matrix and performed the normalization in Chicdiff with the following setting parameters: norm="fullmean", score=3, RUexpand=3L. Di-tag reads between different bait regions were removed from the analysis.

For the classification of enhancer-interacting regions in Extended Data Fig. 1d, we used promoter annotations from the latest version of Ensembl Regulatory Build^91^, CTCF binding sites at E12.5 from publicly available data (GSE181383)^77^, putative enhancers based on H3K27ac occupancy (from E10.5 to E12.5) and polycomb associated H3K27me3 marked regions (at E10.5 to E12.5) from the ENCODE database^81^. We further filtered promoters by only keeping those within ±2.5 kb around TSSs that were transcribed (TPM > 0.5 from RNA-seq data in ENCODE database) in at least one of the following embryonic stages: E10.5, E11.5 and E12.5. CTCF sites were divided into two categories based on whether they were within a TAD or at a TAD boundary. Overlap of interaction peaks with promoters, CTCF sites, enhancers and polycomb regions were computed sequentially, which means peaks were assigned to only one category, and by extending the interaction peaks by ±5 kb.

For the E–P interaction analysis in **Fig. 2, 4, 6 and Extended Data Fig. 6, 7**, we focused on 969 E–P interactions in which the enhancer and interacting gene are both active in at least one tissue. To construct a metaplot profile in Fig. 4, interaction frequencies were scaled as follows: (1) the 5’ end (10 kb around the midpoint of baited enhancer) and the 3’ end (10 kb around the midpoint of interacting promoters) were unscaled; (2) the regions between them have been scaled to 100 kb. Light blue shading indicates 95% confidence intervals estimated by non-parametric bootstrapping. *In vivo* enhancer rank used in Extended Data Fig. 6f, 8c is based on a metric that combines the reproducibility, strength and specificity of staining in the structure(s) of interest and was determined by multiple annotators blinded to genotype (1 = worst; 5 = best)^28^.

To perform *k*-means clustering for E–P interactions in Fig. 4a, normalized interaction frequencies were scaled to the max value among 10 tissues, and clustering was performed in R (v4.1.2) with k = 10 and nstart=30. Clusters were ordered using hclust() with the “ward.D” method and visualized using clusterProfiler (v3.0.4) package^92,93^.

For DNA methylation and DNase signal comparison for interacting and skipped genes in Fig. 2 and Extended Data Fig. 4a-c, we counted the read counts ±1 kb around the TSS of each gene for every enhancer-gene interaction. For comparison to H3K27me3 and H3K9me3 regions, we extended the region analyzed to ±2.5 kb of sequence around the TSS of each gene. For CpG island length analyses in Extended Data Fig. 4g, data was downloaded from the UCSC browser (http://genome.ucsc.edu/cgi-bin/hgTrackUi?g=cpgIslandExt). The differences between interacting and skipped genes were calculated by nonparametric Wilcoxon–Mann–Whitney tests except the comparison for fraction of promoters marked with H3K27me3, which is calculated using chi-squared test.

For E–E interaction analysis in Fig. 4 and Extended Data Fig. 8, we overlapped enhancer interactions with H3K27ac peaks in corresponding tissues in E11.5 embryos (signal >5).

### Generation of enhancer knockout and knockin mice

Enhancer knockout mice were created using a modified CRISPR/Cas9 protocol^33,94^. Briefly, pronuclei of FVB mouse zygotes were injected with a mix of Cas9 protein (final concentration of 20 ng/ul, IDT) and sgRNAs targeting enhancer regions (50 ng/ul) (Extended Data Fig. 5). To replace the ZRS with the fragment of the LacZ sequence, we used a previously described strategy^32^. Briefly, pronuclei of FVB mouse zygotes were injected with a Cas9 protein, a donor plasmid (25 ng/ul) containing a fragment of bacterial *lacZ* sequence and homology arms and sgRNA targeting the ZRS region Cas9 protein^32^ (Extended Data Fig. 9). F_0_ mice were genotyped by PCR and Sanger sequencing using the primers in **Supplementary Table 5**.

### *In situ* hybridization

Whole mount in situ hybridization (ISH) was employed as previously described^32^ to detect *Shh* expression in mouse embryos using digoxigenin-labeled antisense riboprobes (**Supplementary Table 5**), *in vitro* synthesized from a linearized plasmid using RNA Labeling Mix (Roche) and T3 RNA polymerase (Roche). Embryos were fixed with 4% paraformaldehyde (PFA), cleansed in PBT (PBS with 0.1% Tween-20), dehydrated through a methanol series and preserved at -20°C in 100% methanol. For ISH, the embryos were rehydrated, bleached with 6% H2O2/PBT for 15 minutes, and treated with 10 mg/ml proteinase K (PK) in PBT for 20 minutes. Post-PK permeabilization, the embryos were incubated in 2 mg/ml glycine in PBT, rinsed twice in PBT, and post-fixed with 0.2% glutaraldehyde/4% PFA in PBT for 20 minutes. Following three PBT washes, the embryos were transferred to pre-hybridization buffer (50% deionized formamide, 5x SSC pH 4.5, 2% Roche Blocking Reagent, 0.1% Tween-20, 0.5% CHAPS, 50 mg/mL yeast RNA, 5 mM EDTA, 50 mg/ml heparin) for an hour at 70°C, which was after replaced by hybridization buffer containing 1 mg/ml DIG-labeled riboprobe for overnight incubation at 70°C with gentle rotation. The following day, post-hybridization washes were performed at 70°C for 5 minutes with increasing 2xSSC pH 4.5 concentrations: starting from 100% pre-hybridization buffer; 75% pre-hybridization buffer/25% 2xSSC; 50% pre-hybridization buffer/50% 2xSSC; 25% pre- hybridization buffer/75% 2xSSC, followed by 2xSCC, 0.1% CHAPS, twice for 30 minutes at 70°C with gentle rotation. The embryos were then treated with 20 mg/ml RNase A in 2x SSC, 0.1% CHAPS for 45 minutes at 37°C, followed by two 10-minute washes in maleic acid buffer (100 mM Maleic acid disodium salt hydrate, 150 mM NaCl, pH 7.5) at room temperature, and two additional 30-minute washes at 70°C. Samples were then extensively washed in TBST (140 mM NaCl, 2.7 mM KCl, 25 mM Tris-HCl, 1% Tween 20, pH 7.5), blocked with 10% lamb serum/TBST for an hour, and incubated overnight at 4°C with Anti-Dig-AP antibody (Roche, 1:5000) in 1% lamb serum. Excess antibody was removed by washing the embryos in TBST (3×5 minutes), followed by five one-hour TBST washes and an overnight TBST incubation at 4°C. The next morning, embryos were balanced in NTMT (100 mM NaCl, 100 mM Tris-HCl, 50 mM MgCl2, 1% Tween-20, pH 9.5) and alkaline phosphatase activity was visualized by incubating in BM purple reagent (Roche) in the dark with gentle agitation. The reaction was stopped with five 10-minute PBT washes. ISH-treated samples were stored long-term in 4% PFA/PBS and imaged with a Flexacam C1 camera mounted on a Leica M125C stereomicroscope.

### RNA-seq data generation and analysis

Dissected tissues were immediately submerged in RNAprotect Tissue Reagent (Qiagen) and stored at -80 @. Multiple samples from the same tissue and genotype were pooled into at least 1 million cells for each of the two replicates. RNA isolation, preparation of RNA library and transcriptome sequencing was conducted by Novogene Co., LTD (Beijing, China). All RNA-seq experiments were performed in biological replicates. Paired-end reads were mapped to the reference genome (mm10) using STAR (v2.7.9a) software with default parameters^95^ and were counted on RefSeq genes by HTSeq^96^. Differential gene expression analysis was performed using DEseq2 (v3.16)^97^. Genes with adjusted *P*-value < 0.05 were considered differentially expressed.

### DNA FISH in mouse embryonic tissues

DNA 3D-FISH was adapted from previously established methods^98–100^. Fosmid clones from the WIBR-1 library were purchased from the BACPAC Resources Center (for coordinates and names, see **Supplementary Table 4**) and isolated using Large-Construct Kit (Qiagen).

Fluorescent probes were generated using the Nick translation DNA labeling system 2.0 (Enzo) with XFD 488-dUTP or Cyanine-3-dUTP (AAT Bioquest). Unincorporated nucleotides were removed using QIAquick PCR Purification Kit (Qiagen). Probe size (50-500 bps) was analyzed by agarose gel electrophoresis and the incorporation rate was assessed on DeNovix DS-11^101^. Probes were then precipitated with 20X Mouse Cot-1 DNA (Invitrogen) and 20X Salmon Sperm DNA (Invitrogen) and resuspended at 100ng/ul in TE buffer.

Tissues (forelimb, forebrain, midbrain and face) were microdissected from E11.5 mouse embryos and dissociated into single-cell suspension through intubation at 37°C in PBS with collagenase. 50ul of cell suspension (at approximately 5×10^5^ cells/ml) was dropped onto Poly-L-Lysine coated slides (Boster Bio) and incubated for 30 mins at 37°C in a humidity chamber. Slides were then incubated in ice-cold PBS and CSK buffer with 0.5% Triton X-100 for 5 mins respectively, and then fixed in 4% PFA for 10 mins. Slides were sequentially dehydrated in 70%, 80% and 100% ethanol, air dried, and then treated with 400µg/ml RNase A (Fisher Scientific) for 30 mins at 37°C in a humidity chamber. Next, slides were washed with PBS before 10 mins of incubation in 0.1N HCL with 0.5% Tween-20 and 5 mins quenching PBS with 0.02% Tween-20. Samples were then denatured in 70% formamide in 2x SSC pH 7.4 at 80°C for 6 minutes and then dehydrated with 70%, 80%, and 100% ethanol sequentially and air dried. 100ng of probes were diluted in 10µl of hybridization buffer and denatured at 80°C for 10 minutes and pre-annealed for 30-90 mins at 37°C. Pre-annealed probes were added to the cells and covered with a coverslip. Hybridization was carried out in a humidity chamber at 37°C for 16-18 hours. On the next day, slides were washed in 50% formamide in 2x SSC for 3 times, 2x SSC for 3 times, and then 0.1x SSC for twice at 37°C. Slides were then air dried and mounted in 8ul of VECTASHIELD Mounting Medium with DAPI (Vector Laboratories)

### Image acquisition and analysis

Images were obtained on a Zeiss LSM900 Airyscan 2 using a 63X oil objective and an Axiocam 503 mono camera. Lasers were set at 405 (DAPI channel, 3.5% power, 800V gain, 0 offset), 488 (488 enhancer probe channel, 4.0% power, 800V gain, 0 offset) and 561 nm (Cy3 promoter probe channel, 4.0% power, 750V gain, 0 offset) laser lines, and emission bandpass at 400/502 nm (DAPI channel), 496/566 nm (488 probe channel), and 560/700 nm (Cy3 probe channel). SR-4Y multiplex acquisition with a scan speed of 8 was used with a pixel time of 0.5µs and pixel size of 0.04µm; pinhole size was set at 0.2 Airy Units. Z-stacks of 10 slices spanning the nucleus (as determined by the DAPI channel) were taken resulting in an average step size of 0.5µm. Images were deconvoluted using ZEN Blue Software (Zeiss) Airyscan 2 to produce 3D images, and the resulting 3D images were analyzed using Imaris software (Oxford Instruments). We used the Spots module (threshold was set automatically by the software) to computationally identify FISH probe foci. Only foci within the DAPI-stained area containing single probe signals were analyzed to eliminate sister chromatids. The centroids of foci were modeled using PSF-elongation along the Z-axis to create elliptical shaped spots. Inter-probe distances were automatically calculated as the distance in 3D between the centroids of the 488 and Cy3 probe foci. The object-to-object statistics module was used to identify the closest Cy3-promoter foci to each 488-enhancer foci and calculate promoter-enhancer distances. Only pairs with a distance <1.5 µm were considered for further analysis.

### Statistics and reproducibility

No prior analyses were usepaged to determine the sample size before the experiment. The embryos that were not at the correct developmental stage were excluded from data collection. For DNA-FISH image analysis only alleles within the DAPI-stained area and with single probe signals were analyzed to eliminate sister chromatids. Inter-probe distances were measured with the closest distance between a pair of probes and only distances <1.5 µm were considered. For the capture Hi-C and RNA-seq experiment, wild-type and knockin/knockout littermates were randomized and identified only by numbers with genotype unknown to the investigator during data collection and sample processing. For each tissue and corresponding probe set for DNA-FISH, random x-y coordinates were selected and a 9×9 tiled image was taken. For RNA-seq, investigators were blinded to animals’ genotypes during sample collection and library preparation for two knockout lines generated in this study. For ISH experiments in knockin embryos, investigators were blinded to animals’ genotypes during tissue collection and in situ hybridization. For capture Hi-C experiments blinding was not performed because all metrics were derived from absolute quantitative measurements without human subjectivity. For DNA-FISH, after manual data exclusions (see above) foci recognition and distance measurement was done by an automated algorithm (IMARIS).

For comparison of interaction frequencies, histone modifications, DNase accessibility, or inter-probe distances for 3D DNA FISH, no assumptions of normality were made, and all tests were performed using nonparametric Wilcoxon–Mann–Whitney test, nonparametric Fisher’s exact test or Chi-square test. Wilcoxon–Mann–Whitney tests were performed in R using the wilcox.test() performed as a two-sided test. Detailed statistical analyses used in the paper are described in the Methods section. Statistical tests were chosen as appropriate for the data types as described.

### Reporting summary

Further information on research design is available in the Nature Portfolio Reporting Summary linked to this article.

## Data Availability

Sequencing data generated in this study are available at the Gene Expression Omnibus repository with the accession number <u>GSE217078</u>. Several mouse embryonic ChIP-seq / DNase-seq / bisulfite-seq / RNA-seq data for different tissues at E11.5 were downloaded from ENCODE (https://www.encodeproject.org/). The CTCF ChIP-seq data datasets used for comparison were downloaded from GEO (https://www.ncbi.nlm.nih.gov/geo/) under accession numbers <u>GSM5501396</u>, <u>GSM5501397</u> and <u>GSM5501398</u>. Enhancer interaction profiles are available at https://www.kvonlab.org/data/echic. 3D DNA-FISH data are provided as tables in Source data.

## Code availability

Public software and packages were used following the developer’s manuals. The custom code used for data analysis has been deposited at GitHub (https://github.com/kvonlab/Chen_et_al_2024) and Zenodo (https://doi.org/10.5281/zenodo.10594800)^102^.

